# Developmental-stage-specific proliferation and retinoblastoma genesis in RB-deficient human but not mouse cone precursors

**DOI:** 10.1101/356527

**Authors:** Hardeep P. Singh, Sijia Wang, Kevin Stachelek, Sunhye Lee, Mark W. Reid, Matthew E. Thornton, Cheryl Mae Craft, Brendan H. Grubbs, David Cobrinik

## Abstract

Most retinoblastomas initiate in response to the inactivation of the *RB1* gene and loss of functional RB protein. The tumors may form without additional genomic changes and develop after a pre-malignant retinoma phase. Despite this seemingly straightforward etiology, mouse models have not recapitulated the genetic, cellular, and stage-specific features of human retinoblastoma genesis. For example, whereas human retinoblastomas appear to derive from cone photoreceptor precursors, current mouse models develop tumors that derive from other retinal cell types. To investigate the basis of the human cone-specific oncogenesis, we compared developmental-stage-specific cone precursor responses to RB loss in human and murine retina cultures and in cone-specific *Rb1* knockout mice. We report that RB-depleted maturing (ARR3+) but not immature (ARR3-) human cone precursors enter the cell cycle, proliferate, and form retinoblastoma-like lesions characterized by Flexner-Wintersteiner rosettes, then form low or non-proliferative pre-malignant retinoma-like lesions with fleurettes and high p16^INK4A^ and p130 expression, and finally form highly proliferative retinoblastoma-like masses. In contrast, in murine retina, only RB-depleted *immature* (Arr3-) cone precursors entered the cell cycle and they failed to progress from S to M phase. Moreover, whereas the intrinsically highly expressed MDM2 and MYCN contribute to RB-depleted maturing (ARR3+) human cone precursor proliferation, ectopic MDM2 and Mycn promoted only *immature* (Arr3-) murine cone precursor cell cycle entry. These findings demonstrate that developmental-stage-specific as well as species- and cell-type-specific features sensitize to *RB1* inactivation and reveal the human cone precursors’ capacity to model retinoblastoma initiation, proliferation, pre-malignant arrest, and tumor growth.

**Significance Statement:** Retinoblastoma is a childhood tumor that forms in response to mutations in the *RB1* gene and loss of functional RB protein. Prior studies suggested that retinoblastomas arise from cone photoreceptor precursors, whereas mouse models yield tumors deriving from other retinal cell types and lacking human retinoblastoma features. Here, we show that in cultured human retinae, retinoblastomas initiate from RB-depleted cone precursors that are in a specific maturation state and form pre-malignant “retinomas” prior to retinoblastoma lesions, as is believed to occur in retinoblastoma patients. In contrast, Rb-deficient mouse cone precursors of similar maturation state and supplemented with human-cone-precursor-specific oncoproteins fail to proliferate. Thus, human species-specific developmental features underlie retinoblastomagenesis and may challenge the production of accurate mouse retinoblastoma models.

## Introduction

Retinoblastoma is a pediatric retinal cancer that initiates in response to biallelic inactivation of *RB1* or *MYCN* amplification (*MYCNA*) in a susceptible retinal cell type (1). RB1-null retinoblastomas are far more common than the *MYCNA* variety and form either with two somatic *RB1* mutations or with one germline and one somatically inactivated *RB1* allele. Loss of functional RB protein in the cell-of-origin is thought to enable limited proliferation leading to pre-malignant retinomas, from which rare cells escape to form retinoblastoma tumors (2). Retinomas are found at the base of most retinoblastomas, display differentiated histology, and express the senescence markers p16^INK4A^ and p130. The mechanism of escape from the retinoma stage is currently unknown, but likely does not require further mutations since early tumors may lack genomic changes beyond biallelic *RB1* loss (3, 4). Accurate modeling of retinoma formation and escape may provide opportunities to improve treatment and prevent tumorigenesis in genetically predisposed children. However, genetically engineered animal models have not recapitulated the genetic, cellular, and stage-specific features of human retinoma or retinoblastoma genesis.

In humans, germline *RB1* null mutations predispose to retinoblastoma with ~90% penetrance (5). In contrast, mice with germline *Rb1* mutations (6–8) or chimeric biallelic *Rb1* loss (9, 10) fail to develop retinal tumors. Mouse retinal tumors can form when biallelic *Rb1* loss is combined with inactivation of *Rbl1* (encoding p107) (11–14), *Rbl2* (encoding p130) (15, 16), or *Cdkn1b* (encoding p27) (17), or when combined with Mycn overexpression ((18), reviewed in (19)). However, in contrast to the efficient development of human retinoblastoma from rare *RB1*-/- cells, murine retinal tumors formed only when millions of retinal cells had *Rb1* loss combined with one of the other genetically engineered changes, suggesting that still further genomic changes were needed (16). Moreover, although the mouse tumors formed Homer-Wright rosettes, which are common to diverse neuronal tumors (20), they lacked retinoblastoma-specific Flexner-Wintersteiner rosettes and did not progress through a retinoma stage. Furthermore, the mouse tumors prominently expressed retinal interneuron-specific proteins (11–15), whereas human retinoblastomas predominantly express cone photoreceptor proteins (21–25). Specifically, in a series of 40 retinoblastomas, the vast majority of RB-deficient cells expressed protein markers of cones but not other retinal cell types (25). Thus, current mouse models lack the cone protein expression profile, histologic features, and tumorigenesis stages of RB-deficient human retinoblastomas.

The different phenotypes of RB-deficient human retinoblastomas and mouse retinal tumors likely reflect their origins from different retinal cell types. Whereas Rb-deficient mouse retinal tumors were proposed to originate from retinal progenitor cells (14), from horizontal or amacrine interneurons (11, 26), or from a multipotent cell capable of producing amacrine and horizontal cells (16), human retinoblastomas appear to develop from the normally post-mitotic cone precursors that are poised to become mature cone photoreceptors (27, 28). In keeping with the human tumors’ cone-dominant protein profile, RB depletion enabled proliferation of prospectively isolated human cone precursors but not other retinal cells, and proliferation depended on transcription factors (RXRγ and TRβ2) and oncoproteins (MDM2 and MYCN) that are highly expressed during human cone maturation (25, 28). The cone precursors responding to RB loss also expressed maturation markers such as L/M-Opsin and cone arrestin (ARR3) and formed tumors with retinoblastoma-related histology, protein expression, and ultrastructure in orthotopic xenografts (28). In contrast, mouse models with photoreceptor-specific inactivation of *Rb1* or combined inactivation of *Rb1*, *Rbl1*, and *Tp53* failed to produce cone-derived retinoblastomas (29), although *Rb1* loss at the retinal progenitor cell stage permitted subsequent cone precursor cell cycle entry (11). These findings suggest that *Rb1-null* mouse cone precursors lack features that enable extensive proliferation and tumorigenesis in their human counterparts.

To probe the basis for the human but not mouse cone precursors’ proliferative response to RB loss, the current study aimed to identify the developmental stage at which human RB-deficient cone precursors proliferate, to characterize murine Rb-depleted cone precursor responses at the equivalent stage, and to determine whether ectopic expression of oncoproteins (MDM2 and MYCN) that are intrinsically highly expressed in human but not mouse cone precursors enables Rb-depleted mouse cone precursor proliferation. The study also examined the longer-term behavior of RB-depleted cone precursors in human retina cultures. Our findings reveal that RB-depleted maturing human but not mouse cone precursors recapitulate important events in human retinoblastomagenesis and that the cone-related MDM2 and Mycn fail to sensitize maturing murine cones to RB deficiency. These findings provide a novel setting in which to explore retinoblastomagenesis mechanisms and suggest that fundamental differences in murine and human cone precursors may challenge the development of ontogenically accurate retinoblastoma models.

## Results

### RB depletion induces cell cycle entry solely in maturing (ARR3+) human cone precursors

As prior studies suggested that retinoblastomas originate from cone precursors (28), we aimed to define the human cone precursor maturation stage that is sensitive to RB loss. Towards this end, human retinae were cultured on a membrane (30) that largely preserves retinal structure and cellcell interactions (Fig. 1A). With this system, all cell types survived for at least three weeks except Brn3+ retinal ganglion cells, which died within 7 days in culture (DIC) likely due to optic nerve transection (31). As the retina develops in a central-to-peripheral gradient (32, 33), these cultures allowed us to assess the effects of RB loss at different cone precursor maturation states as revealed by topographic position and expression of cone maturation markers (Fig. 1A). As a proxy for the onset of cone maturation we immunostained for ARR3, whose initial expression was found to coincide with the emergence of cone outer segments and the appearance of apically positioned highly concentrated actin filaments that are implicated in outer segment development (Fig. S1).

**Figure 1.**
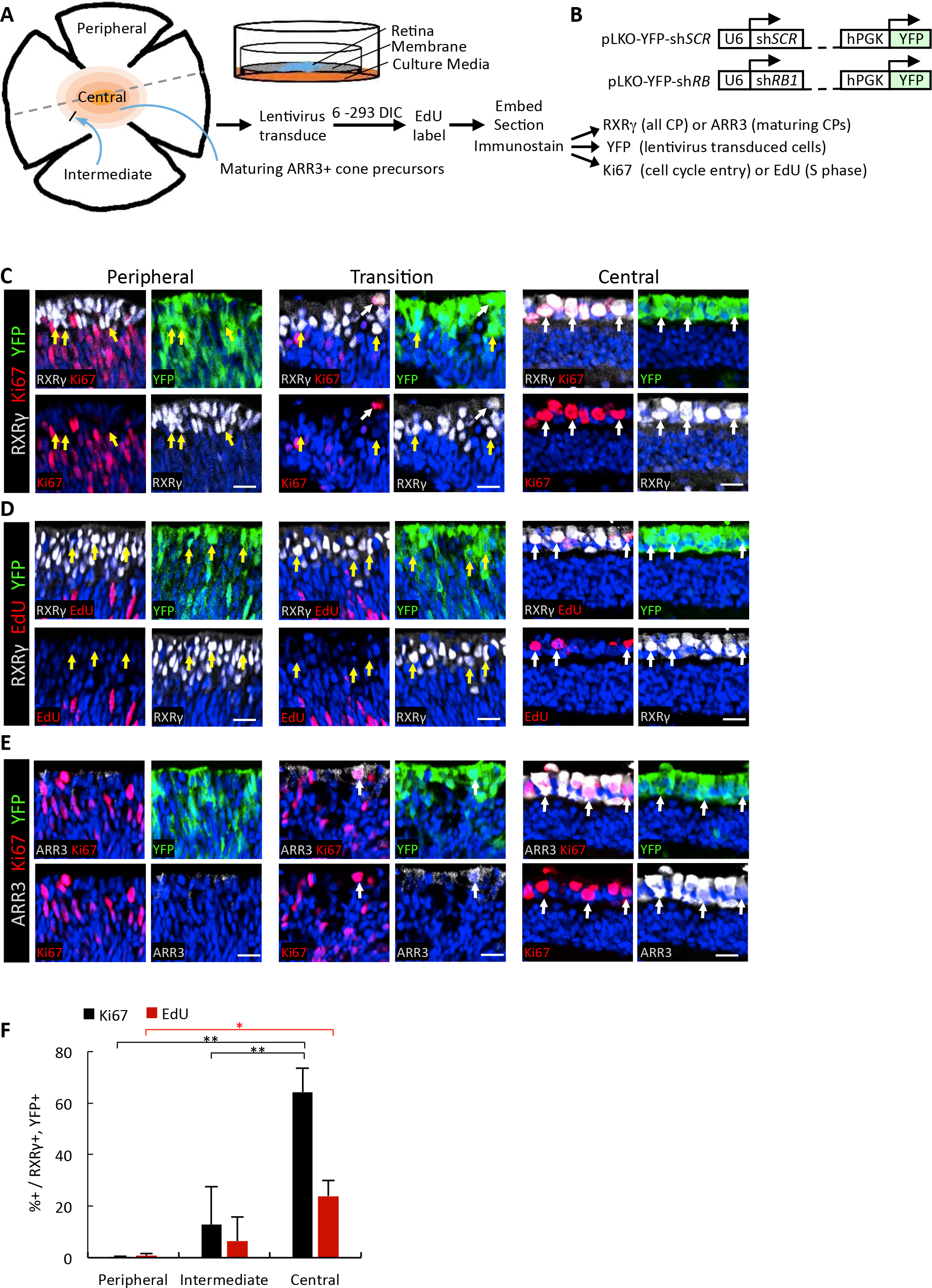
RB depletion induces cell cycle entry in maturing (ARR3+) human cone precursors. **A.** Explanted retinae were cultured on membrane, transduced with lentiviral sh*RB1* or sh*SCR* control shRNA vectors, and EdU-labeled for 4 h immediately prior to harvest. Shading in the intermediate and central retina depicts progressively higher maturation of ARR3+ cone precursors. Dashed line represents a section across the cultured retina that may be used to compare cone precursor behaviors according to topographic position. **B.** Lentiviral constructs used for intact retina transductions. **C-E.** Maturation-related cell cycle entry at 12 days post-RB KD in week 18 human retina examined in peripheral or central retina (as in panel 1A) or in a transition zone displaying the most peripheral ARR3+ cells. Each image is divided into four panels to assess cone markers (RXRβ or ARR3, white), cell cycle markers (Ki67 or EdU, red), lentivirus transduced cells (YFP, green) and DAPI-stained nuclei (blue). Scale bars, 20 μm. **F.** Percentages of Ki67+ or EdU+ cells (stained separately in adjacent sections) among RXRγ+,YFP+ cone precursors in peripheral, intermediate, and central retina. Error bars, standard deviation (SD). Significance assessed by t-test (*, p<0.01, **, p<0.001).

To assess the effects of RB loss, retinae were transduced with lentivirus that co-expressed yellow fluorescent protein (YFP) and either a validated *RB1*-directed short hairpin RNA (shRNA) (pLKO-YFP-*shR8733*) or a non-targeting “scrambled” shRNA control (pLKO-YFP-sh*SCR*) (28) (Fig. 1B). After 12 DIC, retinae were EdU-labeled, sectioned, stained either for RXRy to identify all cone precursors (34, 35) or for ARR3 to identify maturing cone precursors; for YFP to identify transduced cells; and either for Ki67 to detect cell cycle entry or for EdU to monitor S phase entry (Fig. 1A). In retinae explanted and sh*RB1*-transduced at week 18 and examined at 12 DIC, YFP was detected throughout the retina, indicative of transduction of all retinal layers (Fig. S2). However, as YFP was strongest in the cone-rich outer nuclear layer (ONL), imaging was optimized to resolve YFP+ cells in these regions.

In the more mature central retina, most if not all YFP+,RXRγ+ cone precursors were Ki67+ and many incorporated EdU (Fig. 1C, D, F). All Ki67+ and EdU+ cells in this region were strongly ARR3+ (Fig. 1E). In contrast, YFP+,RXRγ+ cone precursors in the immature peripheral retina lacked ARR3 and failed to express Ki67 or incorporate EdU (Fig. 1C-F). At intermediate positions where cone precursors weakly expressed ARR3, a low proportion of YFP+,ARR3^weak^ cells entered the cell cycle and no YFP+,ARR3(-) cells entered the cell cycle (Fig. 1E, F). Retinae transduced with the sh*SCR* control had no Ki67 or EdU incorporation in YFP+,RXRγ+ cone precursors at any position, although sh*SCR*-transduced as well as sh*RB1*-transduced retinae had the expected Ki67+,RXRγ− retinal progenitor cells or glia (28) in the peripheral retina neuroblastic layer (Fig. S3). Thus, in intact human retina, RB-depleted cone precursors acquired the ability to enter the cell cycle at a maturation stage coinciding with the onset of ARR3 expression.

### Rb loss induces cell cycle entry in immature (Arr3-) but not in maturing (Arr3+) murine cone precursors

We next asked whether Rb loss enables murine cone precursor proliferation at the same developmental stage as in cultured human retina. To examine effects of Rb loss on murine cone precursors *in vivo*, we produced mice carrying Cre-dependent *Rb1*^*lox*^ alleles (36) and expressing Cre recombinase under control of the cone-specific *red green pigment* (*RGP*, also known as *Opnllw*) promoter (37). The *RGP-Cre* strain was chosen because the *RGP* promoter is active in mouse cones beginning at ~ postnatal day (P) 8, soon after the age when murine Arr3 protein is first detected (38, 39). At both P10 and P28, Rb was readily detected in Rxrγ+ or Arr3+ cone cells in control *Rb1*^*lox/lox*^ mice but not in *RGP-Cre;Rb1^lox/lox^* littermates (Fig. 2A and Fig. S4, white arrows). In contrast, Rb was equally prominent in inner nuclear layer (INL) cells of both genotypes (Fig. 2A and S4, yellow arrows). Thus, *RGP-Cre*;*Rb1*^*lox/lox*^ mice were expected to reveal effects of cone-precursor-specific Rb loss beginning at or before P10, soon after the onset of Arr3 expression. However, no Ki67 signal was detected among 2221 Arr3+ or L/M-Opsin+ cones examined at P8, P10, P15, and P28 (Fig. 2B). Furthermore, none of 50 *RGP-Cre*;*Rb1*^*lox/lox*^ mice formed retinal tumors after aging for 6-12 months, and no retinomas or other hyperplasias were detected among three histologically examined retinae. Thus, *Rb1* knockout in maturing murine cone precursors failed to induce cell cycle entry or tumorigenesis.

**Figure 2.**
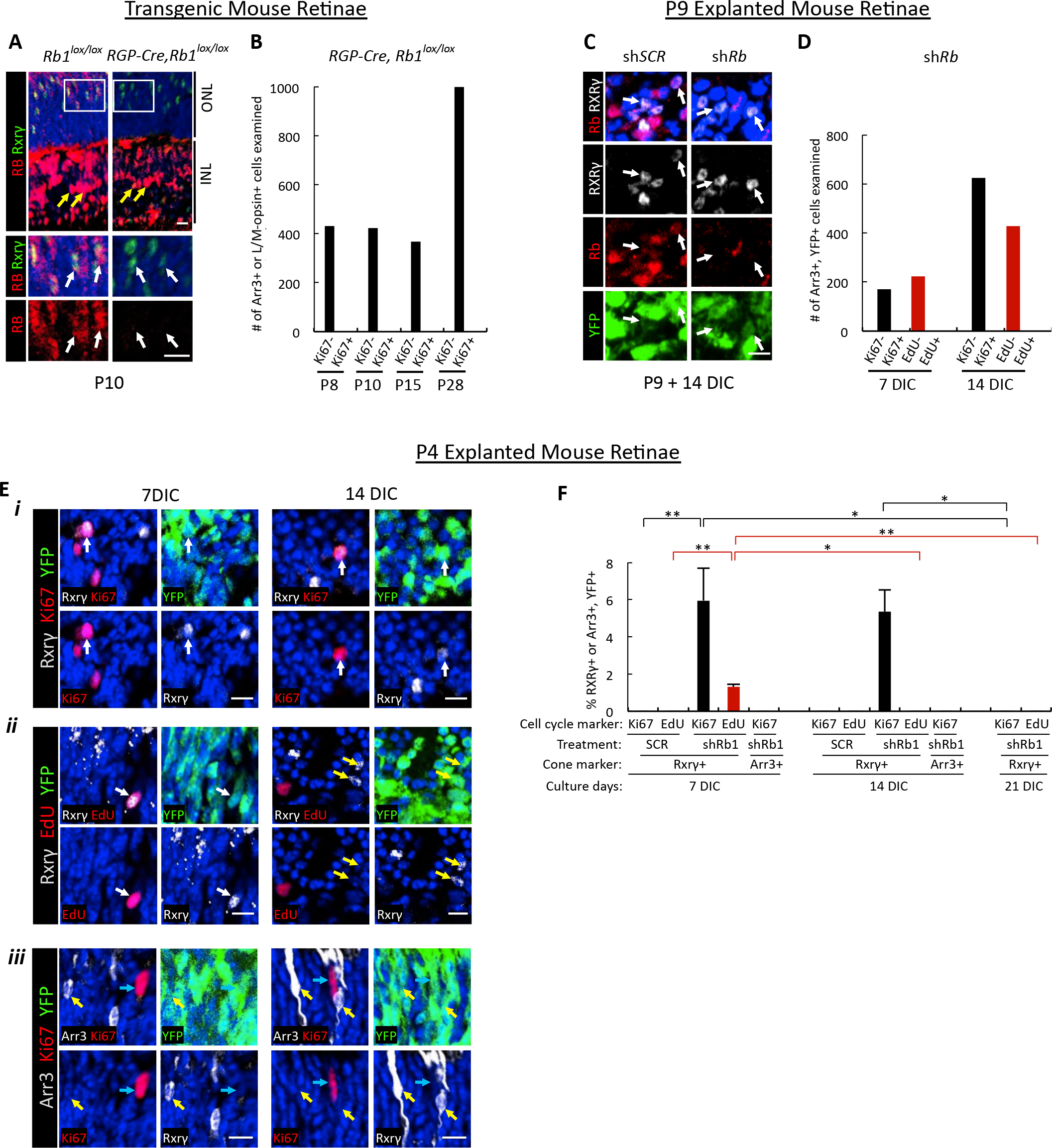
*Rb1* loss induces cell cycle entry in immature (Arr3-) murine cone precursors. **A.** Cone-specific *Rb1* knockout in *RGP-Cre;Rb1*^*lox/lox*^ but not in littermate *Rb1*^*lox/lox*^ mouse retinae at postnatal day (P) 10. Upper panels show sections traversing the outer nuclear layer (ONL) and inner nuclear layer (INL). Boxed ONL regions are enlarged in lower panels. White arrows show Rxrγ+ cones and yellow arrows show Müller glia. **B.** Number of Arr3+ or L/M-Opsin+ cells analyzed for Ki67 expression at various ages in *RGP-Cre,Rb1*^*lox/lox*^ retinae. **C.** P9 explanted retinae transduced with pLKO-sh*Rb1* or pLKO-*SCR* control lentivirus stained for Rb, Rxrγ and YFP at 14 DIC. Arrows show Rb staining in sh*SCR*-transduced but not in sh*Rb1*-transduced YFP+,Rxry+ cone precursors. **D.** Number of Arr3+,YFP+ cells analyzed for Ki67 expression and EdU incorporation at 7 and 14 days post Rb KD. **E.** P4-explanted and sh*RB1*-transduced mouse retina co-immunostained for cone markers Rxry or Arr3 (white), sh*Rb1*-transduction marker YFP (green), and proliferation markers Ki67 or EdU (red). Examples of ONL Rxrγ+,YFP+,Ki67+ (i), Rxrβ+,YFP+,EdU+ (ii), and Arr3+,YFP+,Ki67- (iii) cells at 7 and 14 DIC are shown. White arrows, proliferation-marker-positive cone precursors; yellow arrows, proliferation-marker-negative cone precursors; blue arrows, proliferation-marker-positive Arr3-negative cells. **F.** Quantitation of YFP+,Rxrβ+ or YFP+,Arr3+ cells co-stained for Ki67 or EdU from explanted P4 retinae transduced with sh*Rb1* or sh*SCR* and analyzed at 7 DIC, 14 DIC, and 21 DIC. Error bars, SD. Significance was assessed by ANOVA with Tukey HSD Post-hoc Test. (*, p<0.05,**, p<0.001). Scale bars in A, C, E, 10 μm.

The lack of cone precursor cell cycle entry in *RGP-Cre;Rb1*^*lox/lox*^ mice contrasted with the response of maturing (ARR3+) cone precursors in cultured human retinae. To determine whether maturing murine cone precursors can enter the cell cycle when Rb is depleted *ex vivo*, under the same conditions as used to examine human retinae, a validated murine *Rb1*-directed shRNA (40) was delivered into explanted murine retinae using the same YFP-marked lentiviral vector as used for human retina analyses. We first examined effects of Rb knockdown (KD) on murine retinae explanted at P9, similar to the age of *Rb1* knockout in *RGP-Cre;Rb1*^*lox/lox*^ mice and after onset of Arr3 expression. sh*Rb1* transduction of P9 retinae was confirmed to deplete Rb protein in cones and other cells (Fig. 2C), but did not induce Ki67 or EdU incorporation in Arr3+,YFP+ cells at 7 or 14 DIC (Fig. 2D) (Ki67: 0/796 cells examined; EdU: 0/650 cells examined). Thus, Rb depletion failed to induce cell cycle entry in maturing Arr3+ murine cone precursors under conditions where ARR3+ human cone precursors robustly entered the cell cycle.

The Rb-deficient Arr3+ murine cone precursors’ inability to enter the cell cycle both *in vivo* (after *RGP-Cre*-mediated disruption at ~P8) and *in vitro* (after *shRb1* transduction at P9) contrasted with the prior detection of Ki67+ cone precursors at P8 after *Rb1* knockout in retinal progenitor cells (11). In that study, the Ki67+ cone precursors observed at P8 most likely lacked Rb from the time when they were born (between embryonic days (E) 10-16 (41)) and were immature (Arr3-). To determine if Rb-depletion of immature murine cone precursors can elicit cell cycle entry, murine retinae were explanted and transduced at P4, which is ~ six days after cone genesis is completed (41) and ~ two days before the onset of Arr3 expression ((38) and Fig. S1). Indeed, Rb KD in P4 explants induced Ki67 expression and EdU incorporation in Rxrγ+ cones at 7 DIC (Fig. 2E, F). However, we detected Ki67+ but not EdU+ cones at 14 DIC and neither Ki67+ nor EdU+ cones at 21 DIC (Fig. 2F). As we did not detect cleaved caspase 3+, Rxrγ+ cells (data not shown), the Rb-depleted cone precursors most likely had not died but exited the cell cycle, consistent with the presence of surviving *Rb1*−/− cone precursors *in vivo* (11).

Notably, in P4 explanted retina, Arr3 was induced in a subset of cone precursors after 7 and 14 DIC, indicative of maturation *in vitro*. However, whereas ~ 5% of Rxrγ+ cone precursors were Ki67+ at 7 and 14 DIC, no Arr3+ cone precursors were Ki67+ at 7 or 14 DIC (Fig. 2Eiii and 2F) (0/517 cells examined at 7 DIC (p=0.040) and 0/598 cells examined at 14 DIC (p=0.019), for comparison of Rxr㬴+ and Arr3+ cells at 7 and 14 DIC, respectively; ANOVA with Tukey HSD post-hoc test). Thus, Rb KD at P4 enabled limited cell cycle entry of post-mitotic immature (Arr3-) but not maturing (Arr3+) murine cone precursors.

### MDM2 and Mycn promote cell cycle entry of immature but not maturing murine cone precursors

The cell cycle entry of RB-depleted maturing (ARR3+) human but not murine cone precursors suggested that maturing murine cone precursors lack proliferation-related circuitry that is present in their human counterparts. As ARR3+ human but not mouse cone precursors prominently express MDM2 and MYCN (25), and as MDM2 and MYCN were critical to the RB-depleted human cone precursors’ proliferation (28), we examined whether increased MDM2 and/or Mycn can enable Rb-deficient maturing murine cone precursor proliferation in our *in vivo* and *in vitro* models.

Since MDM2 promotes MYCN expression in retinoblastoma cell lines (42), we first explored whether ectopic MDM2 induces endogenous Mycn expression and Rb-deficient cone precursor proliferation in the *RGP-Cre*;*Rb1*^*lox/lox*^ background. As MDM2 is initially expressed at the onset of ARR3 expression in the human retina (Fig. 3A), we simulated the human cone precursor MDM2 expression timing using an *RGP-MDM2* transgene (Fig. 3B), which is induced at or soon after the onset of Arr3 expression (38, 39). The transgene expressed the full-length human MDM2-X1 isoform produced from the p53-regulated *MDM2 P2* promoter (43), chosen because full-length MDM2 predominates in cones and retinoblastoma cells ((42) and DL Qi and DC, data not shown) and because the cone-related RXRγ up-regulates the *P2* promoter (25). In each of five transgenic strains, MDM2 was first detected in L/M-Opsin+ or Arr3+ cones at P9 and expression increased at P15 and P20 (Fig. 3C, D), resembling human MDM2 expression timing (Fig. 3A). However, *RGP-MDM2;RGP-Cre*;*Rb1*^*lox/lox*^ retinae showed no higher Mycn compared to wild type controls at P10 and P20 (Fig. S5A), no Ki67+ cells among 1499 Arr3+ or L/M-Opsin+ cells examined at P8, P10, P15 or P20 (Fig. 3E), and no tumors among 62 mice aged for 6 - 12 months. Thus, ectopic MDM2 did not enable maturing *Rb1* null mouse cone precursors to induce Mycn or enter the cell cycle *in vivo*.

**Figure 3.**
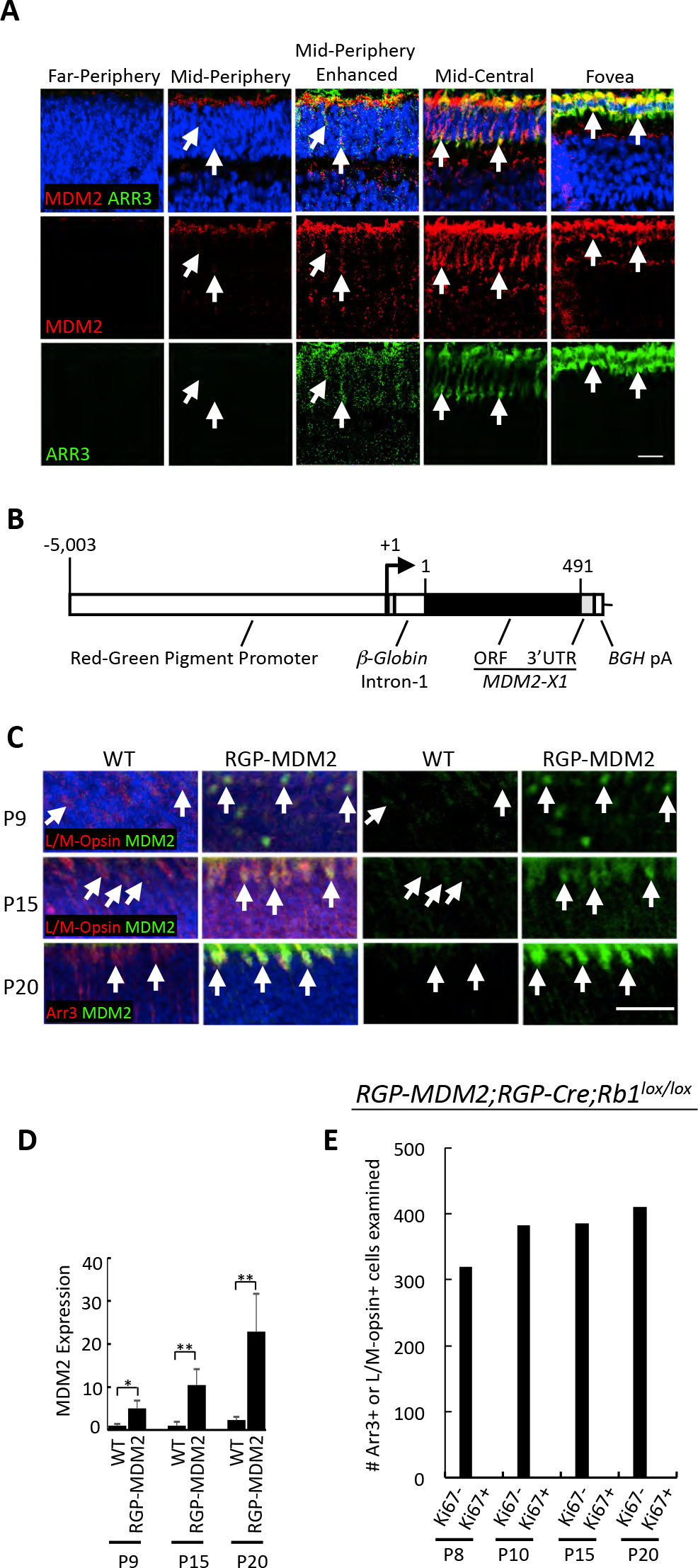
Cone-precursor-specific MDM2 expression fails to sensitize murine cone precursors to *Rb1* knockout *in vivo*. **A.** Coincident onset of MDM2 and ARR3 expression in the developing human retina. Week 21 retina section immunostained for MDM2 (red) and ARR3 (green) and imaged in a peripheral to central series of increasing maturity. Weak MDM2 and ARR3 co-expression was first detected in a mid-peripheral region, which is elucidated by enhancing the same images in the adjacent column. Prominent MDM2 and ARR3 co-expression was observed in mid-central retina and fovea. **B.** *RGP-MDM2* transgene structure with *Opn1lw* promoter, *beta-globin* intron, human *MDM2* isoform X1 open reading frame, and bovine growth hormone poly(A) sequence. Numbers indicate *RGP* promoter base pair position relative to transcription start site and MDM2-X1 amino acid positions. **C.** Increasing expression of human MDM2 (green) in L/M-Opsin+ or Arr3+ (red) cones in *RGP-MDM2* transgenic but not in wild type (WT) mice at P9, P15, and P20. **D.** Quantitation of MDM2 expression in L/M-Opsin+ or Arr3+ cells in *RGP-MDM2* versus WT mice. Error bars, SD. E. Number of Arr3+ or L/M-Opsin+ cells analyzed for Ki67 expression at various ages in *RGP-MDM2;RGP-Cre;Rb1*^*lox/lox*^ retinae. Scale bars, 20 μm. Significance assessed by t-test (*, p<0.05, **, p<0.001).

We next examined the individual and combined effects of ectopic MDM2 and Mycn on Rb-deficient murine cone precursors under the *in vitro* culture conditions that had enabled human cone precursor proliferation (Fig. 4A). In these studies, MDM2 was expressed from the *RGP-MDM2* transgene (Fig. 3B) and appeared *in vitro* with timing similar to that *in vivo* (Fig. S5B). Rb was depleted with pLKO-sh*Rb1* (Fig. 1B), and was confirmed in the context of ectopic MDM2 and Mycn (Fig. S5C). Finally, murine wild type Mycn or a stabilized Mycn protein (Mycn^T58A^) (44) was expressed from the “BN” lentiviral vector (Fig. S5D). Transduction of P9 retinae with BN-Mycn enabled 2- to 6-fold overexpression in ~ 10% of Rxrγ+ cone precursors, whereas transduction of P4 retinae with BN-Mycn^T58A^ enabled 2- to 6-fold Mycn overexpression in ~ 13% of infected Rxrγ+ cone precursors (Fig. S5E, F).

**Figure 4.**
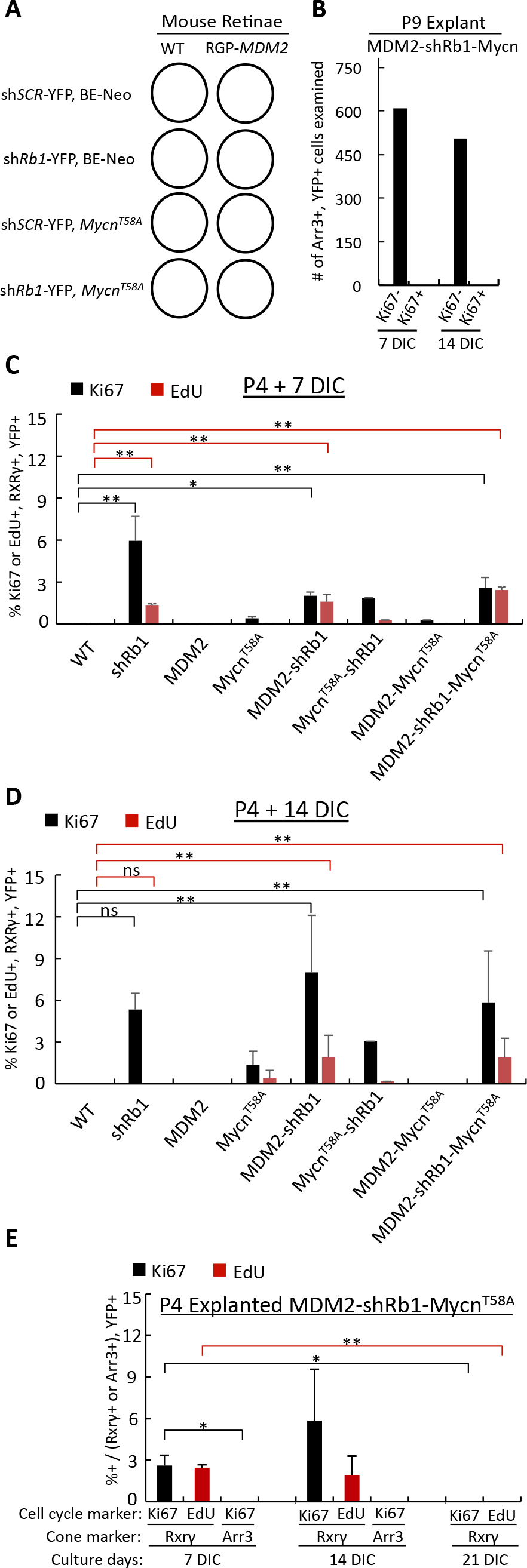
MDM2 and Mycn promote cell cycle entry of immature but not maturing murine cone precursors. **A.** Experimental design for lentiviral transductions of WT or *RGP-MDM2* retinae. All retinae were co-transduced with sh*Rb1* or sh*SCR*, and with BE-Neo-*Mycn*^*T58A*^ or BE-Neo vector. **B.** Number of YFP+, Arr3+ cells from P9 *RGP-MDM2* explanted retinae analyzed for Ki67 at 7 and 14 days post co-transduction with sh*Rb1* and BE-Neo-Mycn^T58A^ **C, D.** Quantitation of YFP+,Rxry+ cells co-stained for Ki67 or EdU in P4 WT or *RGP-MDM2* transgenic retinal explants cultured for 7 or 14 days post-transduction. Labels indicate oncogenic variables (sh*Rb1*, MDM2, Mycn) and omit the applied control vectors or WT genotype. **E.** Quantitation of YFP+,Rxry+ cells or YFP+,Arr3+ cells co-stained with Ki67 or EdU from P4 *RGP-MDM2* retinal explants transduced with sh*Rb1* and BE-Neo-Mycn^T58A^ and analyzed at 7 DIC, 14 DIC, and 21 DIC. Error bars, SD. Significance assessed by ANOVA with Tukey HSD Post-hoc Test (*, p<0.05); **, p<0.001, ns=not significant).

In the *in vitro* setting, transduction of explanted P9 *RGP-MDM2* retinae with sh*Rb1* and ectopic Mycn failed to induce Ki67 at 7 or 14 DIC either when transduced individually (data not shown) or after cotransduction (0/1113 Arr3+,YFP+ cells analyzed, Fig. 4B). Thus, Rb-depleted maturing (Arr3+) murine cone precursors failed to enter the cell cycle regardless of MDM2 and Mycn over-expression.

In contrast, ectopic MDM2 and Mycn promoted cell cycle entry in immature (Arr3-) murine cone precursors after transduction of P4 retinae. Specifically, transgenic MDM2 expression prolonged the Rb-depleted immature cone precursors’ capacity for S phase entry, as Rb KD induced similar proportions of EdU positivity in cone precursors in wild type and *RGP-MDM2* retina at 7 DIC and induced EdU incorporation in *RGP-MDM2* but not wild type retinae at 14 DIC (Fig. 4C, D). Furthermore, transduction of Mycn^T58A^ on its own induced low-level Ki67 expression at 7 DIC and low levels of Ki67 and EdU incorporation at 14 DIC (Fig. 4 C, D). However, co-transduction with *Mycn*^*T58A*^ and sh*Rb1* failed to increase Ki67 expression or EdU incorporation over that induced by sh*Rb1* alone, either in wild type or *RGP-MDM2* retinae (Fig. 4C, D). Similarly co-transduction of *RGP-MDM2* retinae with *Mycn*^*T58A*^ and sh*Rb1* failed to sustain Ki67 expression through 21 DIC or to induce Ki67 expression in Arr3+ cells (Fig. 4E). Thus, in explanted P4 retinae, Mycn^T58A^ induced proliferation in a small proportion of immature cone precursors but did not augment proliferative responses to Rb loss, and failed to enable proliferation of maturing (Arr3+) cone precursors.

### RB-depleted human but not mouse cone precursors proliferate and form retinoma-like and retinoblastoma-like lesions

The above studies indicated that RB-deficient murine and human cone precursors enter the cell cycle at different developmental stages. However, the period of murine cell cycle entry was limited to < 21 days whereas dissociated RB-depleted human cone precursors were reported to remain in the cell cycle *in vitro* for at least 23 days and to form retinoblastoma-like tumors in orthotopic xenografts (28). To investigate this discrepancy, we explored whether RB-depleted murine and human cone precursors in explanted retinae merely enter or also complete the cell cycle. We focused on expression of cyclin B1, which normally accumulates in G2 and M, and on the mitosis-associated phospho-histone H3-Ser10 (pH3-S10). In sh*RB1*-transduced human retinae examined at 12 DIC, RXRyγ+ cone precursors expressed cyclin B1 and pH3-S10 in a central to peripheral gradient (Fig. 5A, C) similar to that of Ki67 and EdU (Fig. 1F). In contrast, sh*RB1*- and Mycn^T58A^-transduced *RGP-MDM2* mouse cone precursors induced neither cyclin B1 nor pH3-S10 (Fig. 5B, Table S1), implying that Rb-depleted and MDM2- and Mycn-overexpressing immature murine cone precursors can enter but fail to complete the cell cycle.

**Figure 5.**
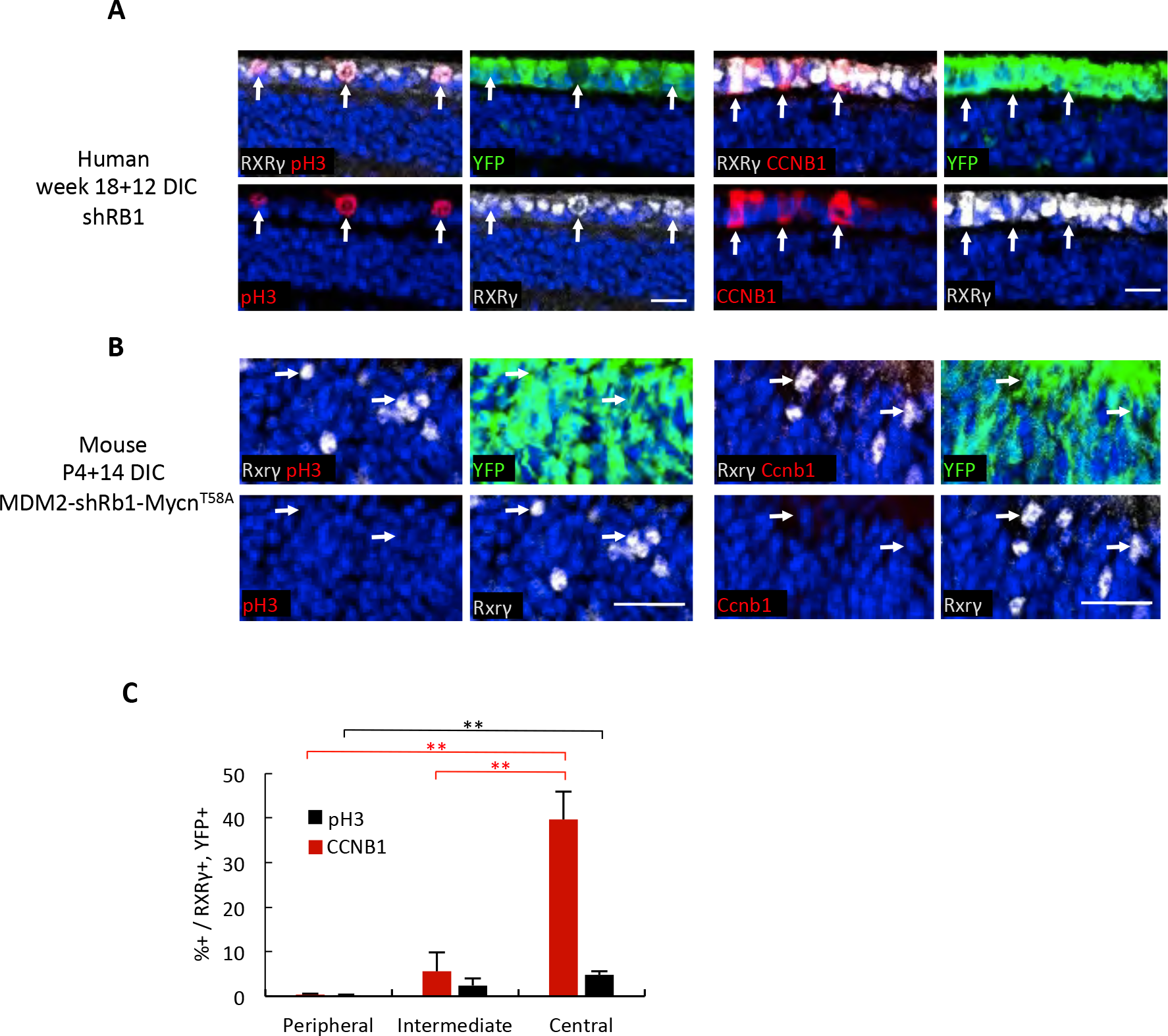
Rb-depleted human but not mouse cone precursors express G2 and M-phase markers. **(A-B)** Immunostaining of G2 and M phase markers cyclin B1 (CCNB1) and phospho-histone H3-Ser10 (pH3) (red) in RXRγ+ (white) cone precursors of 18 week human retina at 12 days post-RB KD (A) but not in Rxrγ+ (white) cone precursors of P4 *RGP-MDM2* mouse retinae at 14 days post-Rb KD and Mycn over-expression (B). Images are from representative regions containing Ki67+,Rxrγ+ and EdU+,Rxrγ+ cells. Scale bars, 20 μm. **C.** Quantitation of % CCNB1+ or pH3+ cells among RXRγ+,YFP+ cone precursors in week 18 human retina at 12 days post *RB1* KD. Serial sections were stained for CCNB1 and pH3 here and for Ki67 and EdU in Fig. 1F. Error bars, SD. Significance assessed by t-test (**, p<0.001).

We next explored the fate of the RB-depleted human cone precursors over extended time in culture. At 30 and 74 DIC, many sh*RB1*-transduced cone precursors, identified by YFP and RXRγ co-expression, were Ki67+ and EdU+ (Fig. 6A-D), although the EdU+ proportion declined relative to that observed at 12 DIC (Fig. 6E). Moreover, the cultured retinae took on a hyperplastic and disorganized appearance, with abundant rosettes, some of which were clearly Flexner-Wintersteiner rosettes, resembling a retinoblastoma tumor (Fig. 6F-H). The Flexner-Wintersteiner rosettes’ ring-like distribution of nuclei was also evident in immunostaining analyses (Fig. 6A-D, yellow circles), which revealed that the rosettes were largely comprised of RXRγ+ cells, many of which were Ki67+ or EdU+. Flexner-Wintersteiner rosettes were not evident in human retinae transduced with control sh*SCR* and cultured in parallel (Fig. 6I), nor in Rb-depleted murine retinae either with or without ectopic MDM2 and Mycn (Fig. S6). Some retinal regions also had Ki67+,RXRγ- cells (i.e., proliferating, non-cone cells); however, some of these were YFP−, indicating that they were not sh*RB1*-transduced (Fig. 6A, B, white arrows), and similar Ki67+ cells were present in retinae transduced in parallel with the sh*SCR* control (Fig. S3 and data not shown), suggesting that they represent proliferating retinal progenitor cells or glia as seen at earlier time points (Fig. S3 and (28)). Thus, in cultured retinae, RB depletion enabled cone precursor cell cycle entry within 12 DIC and enabled proliferation and development of retinoblastoma-like lesions at 30 and 74 DIC.

**Figure 6.**
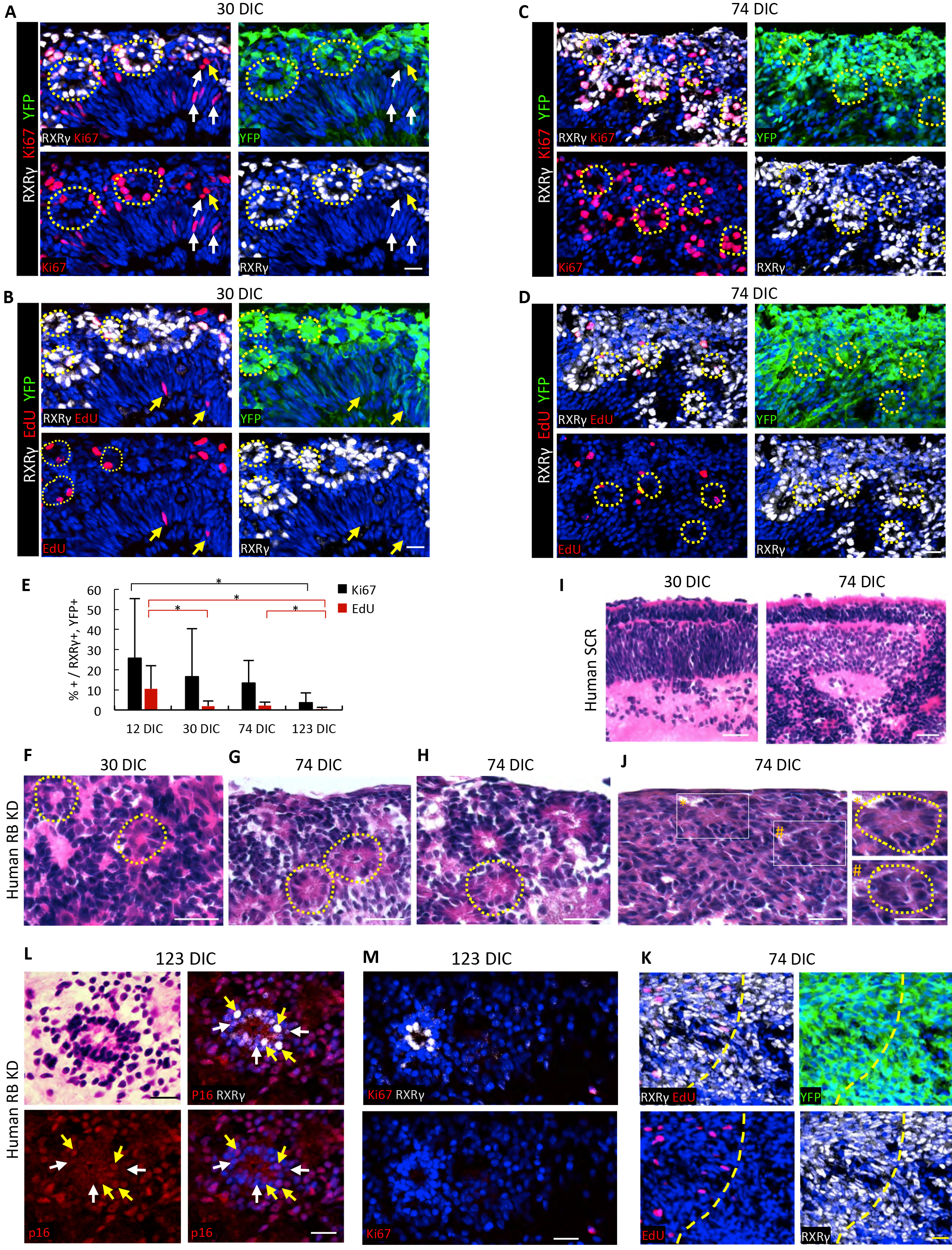
Cone precursor proliferation and retinomagenesis in RB-depleted human but not mouse retinae. **A-D.** Ki67 expression (A, C) and EdU incorporation (B, D) (red) at 30 DIC (A, B) and 74 DIC (C, D) after transduction of week 18 fetal retina. Dashed circles identify ring-like arrangement of RXRγ+ nuclei resembling rosettes. Arrows show proliferating non-cone (RXRγ-) cells, which were either transduced (yellow arrows) or non-transduced (white arrows) and likely represent progenitors or glia. **E.** Quantitation of Ki67 expression and EdU incorporation in RXRγ+ cone precursors in explanted week 18 retinae from 12 to 123 DIC. Data at each age is from central, intermediate and peripheral regions. Error bars, SD. Significance assessed by ANOVA with Tukey HSD Post-hoc Test (*, p<0.05). **F-J**, Hematoxylin and eosin (H&E) staining of human week 18 retina at the indicated DIC after transduction with sh*RB1* (F, G, H, J) or sh*SCR* (I). Dashed circles identify Flexner-Wintersteiner rosettes (F-H) or fleurettes (J, boxed regions at *left* enlarged at *right)*. **K**. Predominance of RXRγ+,YFP+ cells in the same retinoma-like region as shown in panel J. Dashed yellow line separates proliferating (*upper left*) and nonproliferating (*lower right*) regions. **L.** H&E staining (*upper left*) performed after p16 ^INK4A^ immunostaining and imaging of RB-depleted retina at 123 DIC, demonstrating central fleurette with predominantly cytoplasmic p16 in RXRγ^lo^ cells (white arrows) and nuclear p16 in RXRγ^hi^ cells (yellow arrows). **M.** Another representative fleurette at 123 DIC, composed of RXRγ+,Ki67− cells. Scale bars, 20 μm.

At 123 DIC, RXRγ+,YFP+ cells continued to be detected, yet Ki67+ and EdU+ cells were significantly rarer (Fig. 6E). Although the sections analyzed at different DIC were from different retinal regions, the temporal decline in proliferation markers was evident by 74 DIC in regions of disorganized RXRγ+ cells (Fig. 6J, K). Similar features were seen at 119 DIC in an independently transduced retina. Thus, most but not all of the previously highly proliferating RB-depleted cone precursors appear to exit the cell cycle over ~ four months, suggesting that the lesions enter a slowly changing indolent phase.

As the increase in non-proliferating RB-depleted cone precursors was consistent with the production of premalignant retinomas (2), we explored whether cultured retina cone precursor lesions had retinoma-associated features including fleurettes (20) and p16 and p130 expression (2). Indeed, retina transduced with *RB1*-directed shRNA formed fleurettes within highly RXRγ+ regions at 74 DIC (Fig. 6J, K) and formed numerous fleurettes that lacked Ki67 but expressed p16 at 123 DIC (Fig. 6L,M). Moreover, quantitative immunofluorescence imaging revealed that the sh*RB1*-transduced (YFP+) cone precursors at 12 DIC, 30 DIC, and 74 DIC had increased p16 and p130 expression relative to retinae from the fellow eye transduced with sh*SCR* shRNA control (Fig. 7A-D). Notably p16 was mainly nuclear in the RXRγ^hi^ cells that predominated at 12 and 30 DIC and were present at 123 DIC (Fig. 7C, 6L) but was mainly cytoplasmic in the RXRγ^lo^ cells that were more common at 123 DIC (Fig. 6L), as reported for patient-derived retinoma samples (2). In contrast, Rb-depleted murine retinae with ectopic MDM2 and Mycn lacked increased cone precursor p16 or p130 (Fig. 7E-H). Thus, maturing human but not murine cone precursors proliferated and formed retinoma-like lesions in response to RB loss.

**Figure 7.**
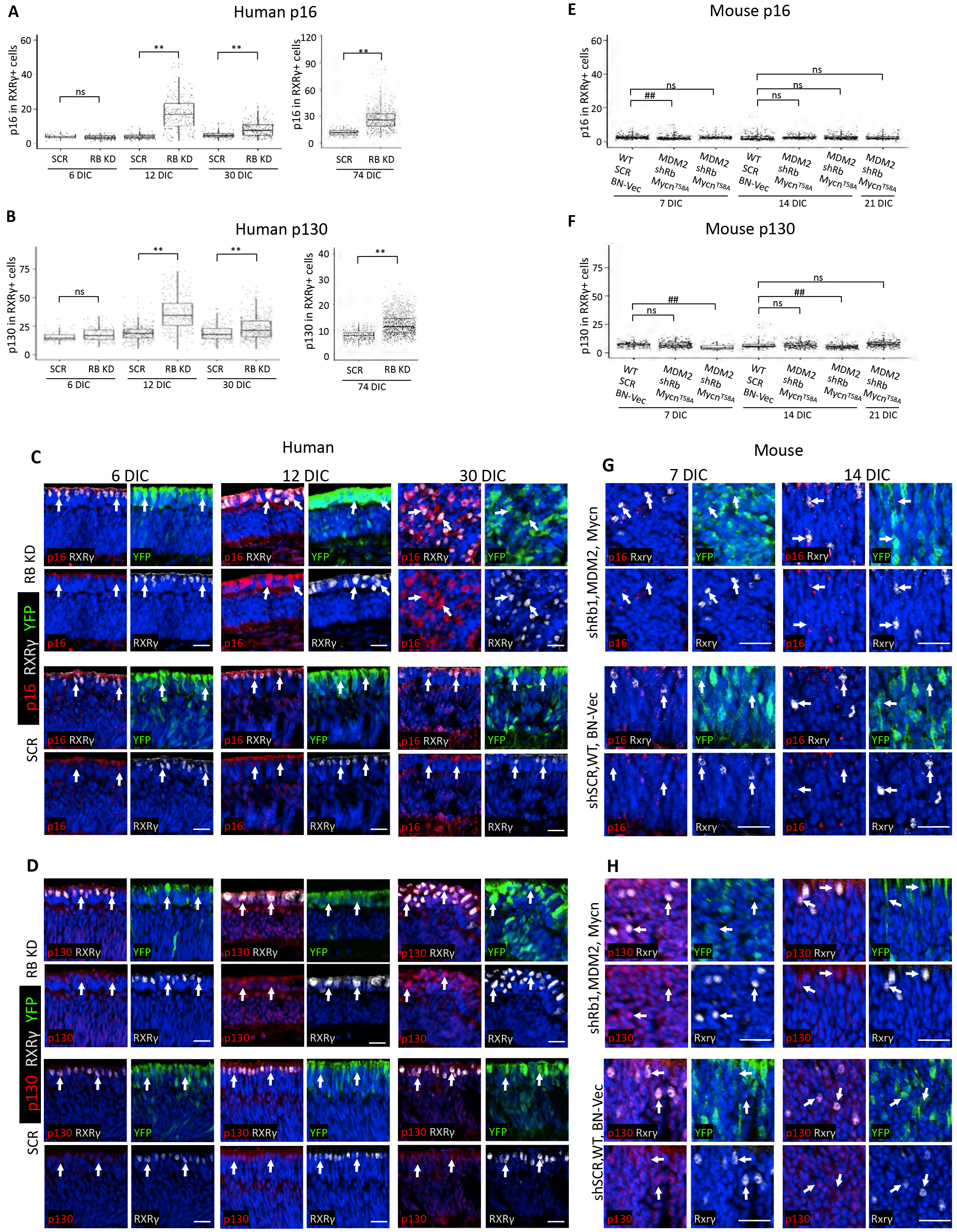
Cone precursor expression of the retinoma-related p16^INK4A^ and p130 in RB-depleted human but not mouse retinae. **A,B,E,F.** Quantitation of p16 and p130 protein in RXRγ+,YFP+ cells in human retina at 6, 12, 30, and 74 days post-transduction with sh*RB1* or sh*SCR* (A,B), and in P4 WT or *RGP-MDM2* mouse retinae at 7, 14, and 21 days posttransduction with sh*SCR*, sh*Rb1*, BN vector, BN-*Mycn*^*T58A*^ (E,F), based on immunostaining shown in panels (C,D and G,H). Each dot represents signal in a quantitatively imaged RXRγ+,YFP+ cell outlined by RXRγ signal. Box plots show median (line inside box), upper and lower quartiles (box borders), and data range (whiskers). Significance assessed by t-test (**,## p>0.0001; ns=not significant). **C,D,G,H.** Immunofluorescence staining of p16 (C, G) and p130 (D, H) in human retina at 6, 12, and 30 days post-transduction with sh*RB1* or sh*SCR* (C,D) and in P4 WT or *RGP-MDM2* mouse retinae at 7 and 14 days post-transduction with (G,H). Scale bars, 20 μm.

The above analyses revealed features of retinoblastoma starting at 30 days post RB KD and features of retinoma at 74, 119, and 123 days post-RB KD. To assess whether RB KD can result in full-blown retinoblastoma, we knocked down RB in week 19 and week 21 retinae and cultured for longer times. In two unrelated retinae, cell masses became visible starting at ~ 8 months post-RB KD and were detected in all quadrants by 9 months post-RB KD (Fig. 8A and data not shown). Histologic examination revealed numerous Flexner-Wintersteiner rosettes (Fig. 8B), consistent with a differentiated retinoblastoma phenotype. The mass was mainly composed of RXRγ+ cells that were in cell cycle as determined by Ki67 expression (Fig. 8C). Interestingly, cytoplasmic p16 expression was evident in proliferating RXRγ^hi^ cells that retained differentiated histology (Fig. 8D), which differed from the nuclear p16 in RXRγ^hi^ cells at earlier time points (Fig. 6L, 7C).

**Figure 8.**
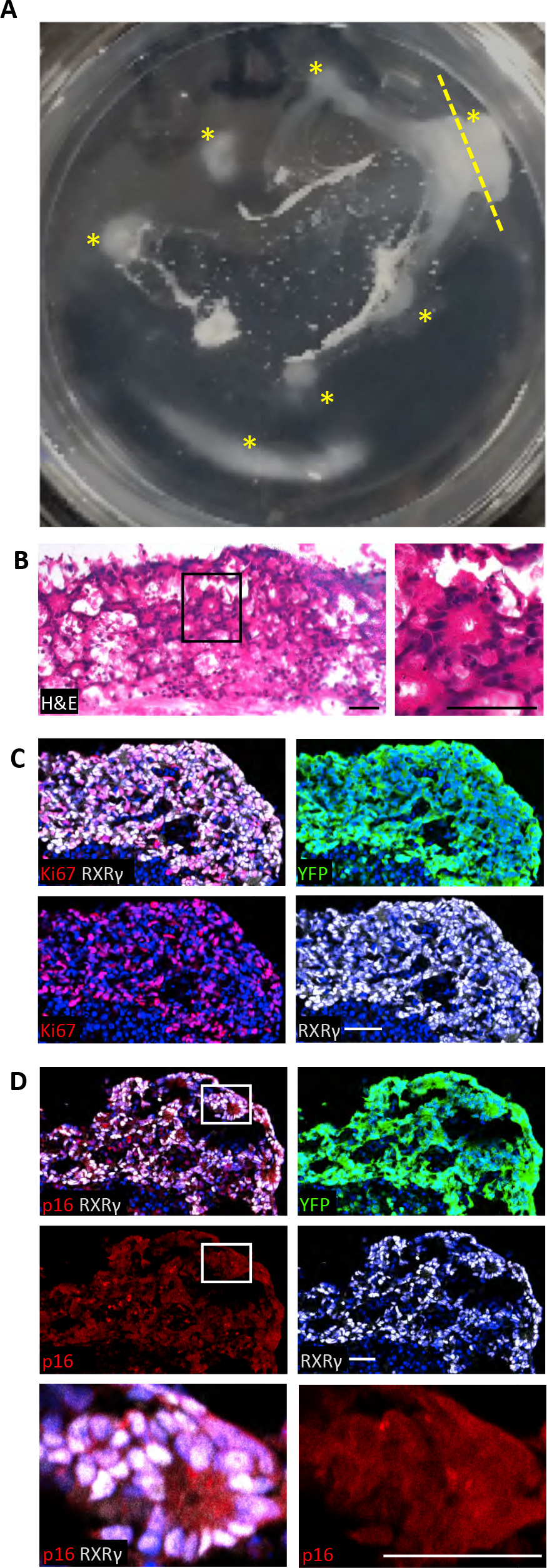
RB-depleted retinal explants form retinoblastoma-like lesions. **A.** Explanted intact retina cultured on a 6-well membrane for 293 days post-RB KD. At least seven distinct cell masses were observed (asterisks). The plane of the section used for H & E and immunostaining is shown as a yellow dashed line **B.** H&E staining reveals retinoblastoma-like histology including Flexner-Wintersteiner rosettes. **C, D.** Ki67 (C) and low-level cytoplasmic p16^INK4A^ (D) are expressed throughout the mass. Boxed region in (D) is enlarged in bottom row. Scale bars, 40 μm.

## Discussion

We report that RB-depleted human *maturing* (ARR3+) cone precursors enter the cell cycle, proliferate, and form retinoma-and retinoblastoma-like lesions, whereas Rb-depleted murine *immature* (Arr3-) cone precursors can enter but not complete the cell cycle (Fig. 9). Maturing Arr3+ murine cone precursors failed to enter the cell cycle both in response to *Rb1* knockout *in vivo* and in response to Rb knockdown initiated either before or after the onset of Arr3+ expression (Fig. 2). Moreover, ectopic MDM2 and Mycn did not enable maturing cone precursor cell cycle entry either *in vivo* or under *in vitro* conditions identical to those that enabled human cone precursor proliferation (Fig. 3, 4). These findings are consistent with past evidence that murine cone precursors fail to proliferate after photoreceptor-specific *Rb1* knockout or combined *Rb1*, *Rb11*, and *Tp53* knockout (29). Thus, maturing (Arr3+) murine cone precursors show an impressive resistance to cell cycle entry, whereas maturing (ARR3+) human cone precursors show an impressive propensity to proliferate under the same conditions.

**Figure 9.**
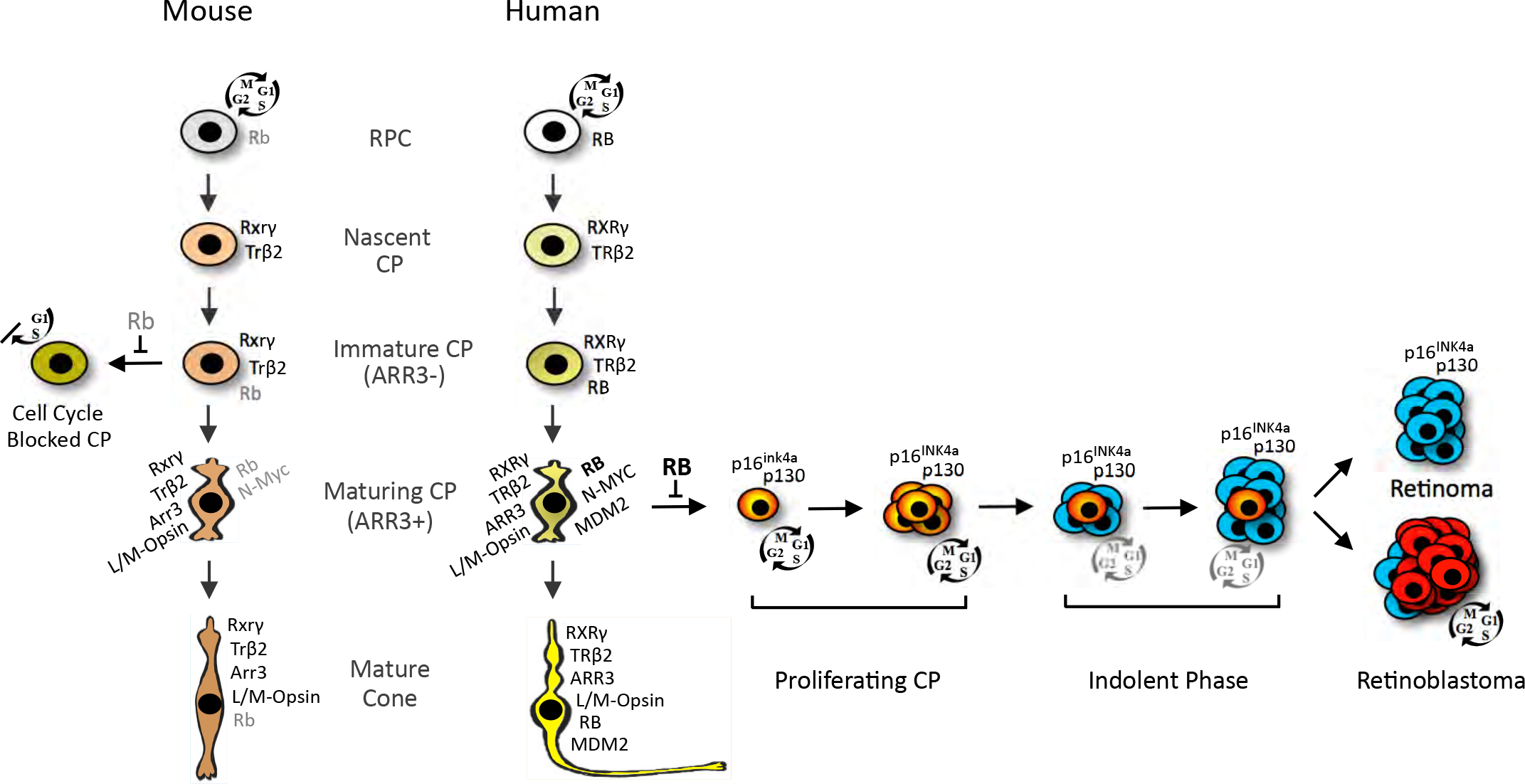
Distinct cone precursor (CP) responses to RB loss in human and murine cone precursors. The cartoon depicts selected protein expression and responses to RB loss in human and mouse nascent CPs, immature (ARR3−) CPs, maturing (ARR3+) CPs, and mature cones. Nascent CPs are RXRγ+, TRβ2+ and have minimal RB expression in humans (34) and mice (45). RB increases during cone precursor maturation in both species but to far higher levels in humans (25, 34, 45). The current work shows that in intact human retinae, RB loss in L/M-Opsin+, ARR3+ cone precursors (with high MDM2, MYCN and RB (25)) elicits cell cycle entry (depicted in *red*), proliferation, and either permanent cell cycle withdrawal (depicted *in blue*) leading to a quiescent retinoma-like state or continued possibly indolent proliferation eventually leading to retinoblastoma. This study also shows that in intact mouse retinae, Rb loss does not induce cell cycle entry in maturing (Arr3+) cone precursors, whereas immature (Arr3-) cone precursors enter but fail to complete the cell cycle, and fail to form retinoma-or retinoblastomalike lesions.

Our findings imply that RB is not needed to enforce cell cycle exit at the time of human cone precursor cell birth, but acquires this role coincident with the onset of ARR3 expression and outer segment growth (Fig. 9). Concordantly, prior analyses indicated that in the human retina, RB is expressed at barely detectable levels in immature cone precursors, and dramatically increases during maturation starting in the fovea at ~ week 14 (reported as post-fertilization week 12, (34)). Rb also was not detected in immature mouse cone precursors from E14 to P5 (45), and was detected at far lower levels during murine versus human cone maturation (25). Thus, the maturation-associated increase in RB expression correlates with RB’s role in suppressing human cone precursor cell cycle entry. If a similar sensitivity pertains *in vivo*, then the first aberrant proliferation of RB-deficient cone precursors may coincide with the onset of ARR3 expression at ~ week 15 in the central retina (46), ~ 6 weeks after the first cone precursors are born at ~ week 8 or 9 (47). A similar interval between cone cell birth and RB-dependent quiescence may apply throughout the retina as cone genesis propagates to the far periphery by week 22 (48).

In this study, the onset of ARR3 expression was used to denote the transition from an “immature” to a “maturing” cone precursor coinciding with cone outer segment budding. While other maturation-related changes may precede ARR3 expression, early outer segment morphogenesis was proposed to reflect the start of a distinct second phase of vertebrate photoreceptor differentiation (49). In this regard human cone precursors may become sensitized to RB loss during this second differentiation phase.

Our finding that RB-depleted maturing (ARR3+) but not immature (ARR3-) human cone precursors enter the cell cycle suggests that in the human retina, ARR3 expression is associated with increased mitogenic signaling. Indeed, cone precursor differentiation is associated with increased MDM2 and MYCN in humans but not mice (Fig. 3 and S5) (25). Although high MDM2 and MYCN were needed for the human cone precursor proliferative response to RB loss (28), ectopic MDM2 and Mycn did not enable cell cycle entry in Rb-deficient maturing murine cone precursors (Fig. 4). As a caveat, the ectopic Mycn might not have been over-expressed to a sufficient level to enable maturing cone precursor proliferation in these experiments. However, maturing mouse cone precursors were resistant to ectopic Mycn overexpression as compared to maturing human cone precursors transduced with the same methods (H.P. Singh, data not shown), suggesting that additional potentially mitogenic signals enable high-level MYCN expression during human but not murine cone maturation. Indeed, maturing human cone precursors express a larger proliferation-related program, beyond high-level MDM2 and MYCN expression (50), that could contribute to the proliferative response to RB loss.

While maturing murine cone precursors did not acquire proliferative potential, prior studies suggested that another murine post-mitotic cell type - horizontal cells - can proliferate and form tumors in response to combined biallelic loss of *Rb1*, biallelic loss of *Rbl2* (encoding p130) and monoallelic loss of *Rbl1* (p107) (26). Like maturing human cone precursors, murine Prox1+ horizontal and amacrine cells have intrinsically high Mdm2 (25), possibly reflecting a similar tumor-prone state. Thus, although the engineered genomic changes were far more extensive, the *Rb1*−/−;*Rb12*−/−;*Rb11*+/− model as well as other models in which tumors derive from Prox1+ interneurons might simulate an oncogenic process that is analogous, yet not orthologous, to human retinoblastoma genesis.

In contrast to the proliferative response of RB-depleted maturing human cone precursors, only Rb-depleted *immature* (Arr3-) murine cone precursors entered the cell cycle (Fig. 2). Immature murine cone precursor cell cycle entry was prolonged by ectopic MDM2 and was independently induced by Mycn (Fig. 4). However, these responses were weaker than the ~ 60% of human cone precursors that entered the cell cycle in central retina (Fig. 1F) and might not be relevant to the development of RB-deficient human retinoblastoma given that the human tumors arise from a distinct developmental stage. The Rb-depleted and MDM2- and Mycn-overexpressing immature murine cone precursors’ failure to express cyclin B1 or pH3-S10 (Fig. 5) indicated that they could enter but could not complete the cell cycle. Rb-deficient murine hepatocytes display similar behavior (51), suggesting that certain Rb-deficient mouse cells abrogate cyclin B1 accumulation as a means to suppress Rb-deficient cell proliferation. However, at present, it is unclear whether human cells are capable of a similar response.

After entering the cell cycle, RB-depleted human cone precursors formed hyperplasias with Flexner-Wintersteiner rosettes and fleurettes, histologic features specific to retinoblastoma and retinoma, respectively (Fig. 6, 7) (2, 20). Flexner-Wintersteiner rosettes were abundant as early as 30 days post-RB KD and cells within the rosettes expressed cone markers at levels similar to the earliest cone precursors that entered the cell cycle, implying that they are the cone precursors’ direct descendants (Fig. 9). Retinoma-like lesions with rare detected at 74, 119, and 123 DIC also expressed cone markers (Fig. 6), suggesting that they derive from Flexner-Wintersteiner-rosette-containing tissue. These lesions had largely cytoplasmic p16, particularly in RXRγ^1^° cells, as seen in patient-derived retinoma regions (2). Finally, multiple masses were observed in a pRB-depleted retina after ~ 9 months. The masses had abundant Flexner-Wintersteiner-rosettes and RXRγ expression similar to that observed at 30 DIC (Fig. 8), resembling differentiated retinoblastoma tissue. The proliferating lesions’ predominantly cytoplasmic p16 in RXRγ^hi^ as well as in RXRγ^1^° cells was a distinct feature of the late-arising retinoblastoma-like versus retinoma-like tissue. Further analyses are needed to assess whether cytoplasmic p16 localization enables the transition from indolent proliferation to differentiated retinoblastoma and whether p16 loss enables a subsequent transition to the de-differentiated retinoblastomas that were reported to have reduced p16 expression (2).

In conclusion, our data supports a model in which human RB-deficient maturing cone precursors initially form proliferative lesions that resemble differentiated retinoblastomas in which the majority of cells exits the cell cycle to form non-proliferating retinomas and a minority remains in the cell cycle, possibly in an indolent state, to form retinoblastoma tumors (Fig. 9). Understanding the parameters that enable proliferating RB-deficient cone precursors to enter or bypass the retinoma state could provide opportunities to suppress retinoblastoma in genetically predisposed children.

## Materials and Methods

### Retinal explant culture

Following informed consent, fetal eyes were obtained from authorized sources with approval by the USC and CHLA Institutional Review Board. The best estimate of the gestational age was determined according to the guidelines put forth by the American College of Obstetrics and Gynecology (52). Post-natal mouse eyes were harvested in retina culture media with the birth day designated P0. Eyes were held on ice for up to 4 h, sterilized in 70% ethanol for 5 seconds, followed by submersion in cold PBS for retina removal. Retinae were harvested as an intact cup, cut radially to flatten, and placed on hydrophilic polytetrafluoroethylene cell culture inserts (Millipore, PICM01250 or PICM0RG50) with photoreceptor side facing down to the membrane. Inserts with retinae were quickly moved to cell culture plates with 1200 μl or 250 μl of Retina Culture Media (IMDM (Corning), 10% fetal bovine serum (FBS; Sigma-Aldrich), Insulin (0.28U/ml, Lilly, USA), β-mercaptoethanol (55 μM, Sigma-Aldrich), Glutamine (Sigma-Aldrich), penicillin, and streptomycin (Corning) added to 6-well or 12-well dishes, respectively. PBS was aliquoted into the unused wells as well as in the spaces between the wells, and the plates incubated at 37 ° C with 5% CO_2_. At various times human (but not mouse) retinae were cut into pie-shaped sectors with each containing central and peripheral regions, and the different sectors were harvested for embedding and analyses.

### Lentiviral transduction of explanted retina

Frozen concentrated virus was thawed on ice and added with FBS, β-mercaptoethanol, Insulin, Glutamine, penicillin, and streptomycin at final concentrations as in retina culture media along with polybrene (4 μg/ml, Sigma-Aldrich, Cat # 107689). The cell culture inserts were moved to empty wells and concentrated virus mix was added on the top of retina. Virus-containing media flowing through the membrane was again placed on the retina every 15-20 minutes. After 3 hours, the cell culture inserts were returned to the wells with fresh media. Half of the media was changed every two days. Retinae were incubated with EdU (10U/ml, Invitrogen, Cat# C10338) for 4 hours prior to harvest and then fixed in freshly prepared 4% paraformaldehyde in PBS for 15 minutes. After three PBS washes, retinae were equilibrated in sucrose (30% in PBS) for 15 minutes and embedded in sucrose OCT mix (2:1) on dry ice, and the blocks stored at −80 °C. Blocks were sectioned at 10 μm thickness on positively charged slides and slides stored at −80 °C.

### Lentivirus production

Lentiviruses were produced in 15 cm dishes by reverse transfection of 3×10^7^ 293T cells using 20 μg lentiviral vector, 5 μg pVSV-G, 10 μg pMDL, 5 μg pREV in 3 ml of serum-free DMEM-HG (Corning). 120 μl of Polyethylenimine (PEI) (0.6μg/ml, Polysciences Inc., Warrington, USA, Cat# 24765) was mixed with 3 ml of serum-free IMDM and incubated for 20 minutes at room temperature. Cells were then mixed with the PEI mix and plated in DMEM-HG media (Corning) with 10% FBS. At ~ 14 to 16 hours, media was replaced with serum-free UltraCULTURE media (Lonza Inc., Basel Switzerland, Cat# 12-725F) and virus was harvested at 60 hours post-transfection. The supernatant was centrifuged at 3000 x g for 10 minutes at 4° C and filtered with 0.45-micron PES membrane filter (VWR, Cat# 10040-470). Virus was concentrated using tangential flow filtration (53), re-filtered (0.45-micron PVDF filter, Millipore, Cat# SLHV033RB) and stored in small aliquots at −80° C. Virus titer was determined using a p24 ELIZA assay (ZeptoMetrix Corporation, NY, USA, Cat# 0801111).

### Mice

All mouse experiments were approved and performed according to the guidelines of the Institutional Animal Care and Usage Committees of Memorial Sloan-Kettering Cancer Center and The Saban Research Institute of Children’s Hospital Los Angeles. *RGP-Cre* (37) and *Rb1*^*lox*/*lox*^ mice (36) were kindly provided by Anand Swaroop and David MacPherson, respectively. *RGP-Cre* and *Rb1*^*lox*/*lox*^ mice were obtained in a mixed 129/SV and C57BL/6 backgrounds and mated with C57BL/6J mice (Jackson Laboratories) for at least 4 generations prior to breeding with other strains.

#### Construction of RGP-MDM2 mice

The *RGP-MDM2* transgene was produced by cloning the human *MDM2* open reading frame within RefSeq variant XM_005268872.4, which encodes the 491 amino acid MDM2 protein isoform X1 in which translation initiates from the first ATG in exon 2, and flanking vector bovine growth hormone poly(A) site from pcDNA3-MDM2 (kindly provided by Ze’ev Ronai) into the BamH1 and PshAI sites 3’ of the rabbit β-globin splice acceptor region of pRGP-β-globin. pRGP-P-globin was produced by replacing the luciferase gene of pGL3-RGP (a kind gift of M. Akimoto and A. Swaroop) with a 640 bp fragment containing the β-globin intron from pCMV-neo-Bam3. pGL3-RGP contained a 5 kb Red-Green Pigment (*Opn1mw*, or “RGP”) promoter as described (37). The transgene was excised as a MluI-AfeI fragment and used to produce five transgenic mouse lines in the C57BL/6J background in the Memorial Sloan-Kettering Cancer Center Transgenic Core Facility and the intact transgene confirmed by Southern blotting. *RGP-MDM2* mice were maintained as heterozygotes by crossing to wild type C57BL/6J mice and were genotyped by PCR using forward primer 5’-CCTCTGCTAACCATGTTCATGCCT-3’ and reverse primer 5’-TCTTGTTCCGAAGCTGGAAT-3’ in the rabbit *beta-globin* intron and MDM2 open reading frame, respectively. A third primer complementary to murine (but not human) Mdm2 *plus* strand (5’-CTCTCGGATCACCGCGCTTCTCC-3’) was used as PCR reaction internal positive control.

### Lentiviral shRNA and cDNA expression constructs Mycn construct

pLKO lentiviral shRNA vectors were obtained from the TRC library (Open Biosystems) or were cloned into the pLKO vector with modifications described before (28). In addition, the RSV promoter was replaced with CMV promoter to enhance the virus production. We used pLKO.1C-YFP-*shRBL*-733 (28) for human RB KD and pLKO.1C-YFP-sh*RB1* (with targeting sequence 5’-AACGGACGTGTGAACTTATAT-3’ (40)) for mouse *Rb1* KD. pLKO.1C-scrambled (sh*SCR*) expressing a non-targeting control shRNA was Addgene plasmid 1864. The lentiviral cDNA expression vector BE-Neo (or “BN”) was as described (28). BN-*Mycn* was produced by inserting mouse *Mycn* cDNA without UTR sequences between the MluI and XbaI sites of BE-Neo. cDNA encoding stabilized Mycn^T58A^ was a gift from Gregory Shackelford (54) and replaced wild type *Mycn* cDNA using In-Fusion (Clontech, Mountain View, CA, USA).

### Immunofluorescence staining and microscopy

Sections were air dried for several minutes and washed with Tris buffered saline (TBS, Bioland Scientific LLC, Cat# TBS01-02). All washes were with TBS. Sections were treated with EDTA (1mM, pH 8.0) for 5 minutes at room temperature followed by blocking and permeabilization in super block (28) for 1 hour. Primary antibodies were diluted in super block (28) overnight at 4 °C or 1 hour at room temperature, followed by incubation with secondary antibodies for 30 minutes at room temperature. Sections were mounted in Mowiol (Calbiochem, cat#475904, DABCO (Sigma cat#D27802)). For quantitation of protein expression, RXRβ+, ARR3+ cells were outlined and quantitated using ImageJ (55).

### Statistical Analyses

For bivariate analysis of frequency data, Fisher’s exact test was used. For multivariable analyses of mean Ki67 and EdU data (e.g., Fig. 4C-E), ANOVA with Tukey HSD Post-hoc test was used. Other pairwise comparisons of mean percentage data were conducted using Student’s t-test as indicated.

## Acknowledgments

We thank Masayuki Akimoto and Anand Swaroop for *RGP-Cre* mice and pGL3-RGP, David MacPherson for *Rb1*^*lox*/*lox*^ mice, Willie Marks for production of *RGP-MDM2* mice, Esteban Fernandez for help with ImageJ analysis, Ze’ev Ronai for pcDNA3-MDM2, Gregory Shackleford for MYCN^T58A^ cDNA, Melissa L. Wilson (USC Department of Preventive Medicine) and Family Planning Associates for assistance in obtaining fetal tissue, David H. Abramson, Nai-Kong Cheung, and Thomas C. Lee for support, and Jennifer Aparicio and Aaron Nagiel for critical reading of the manuscript. We acknowledge the Transgenic Mouse Core Facility of Memorial Sloan-Kettering Cancer Center and the Stem Cell Analytics Core Facility, Animal Care Core Facility, and Imaging Core Facility of The Saban Research Institute of Children’s Hospital Los Angeles. This study was supported in part by the Elsa U. Pardee Foundation, the Fund for Ophthalmic Knowledge, the Larry and Celia Moh Foundation, the Neonatal Blindness Research Fund, the A.B. Reins Foundation, a Young Investigator’s Grant from Alex’s Lemonade Stand Foundation (to S. Lee.), an unrestricted grant to the USC Department of Ophthalmology from Research to Prevent Blindness, New York, NY, and NIH grants P30CA014089 and R01CA137124 (D. Cobrinik).

---

**Figure S1.**
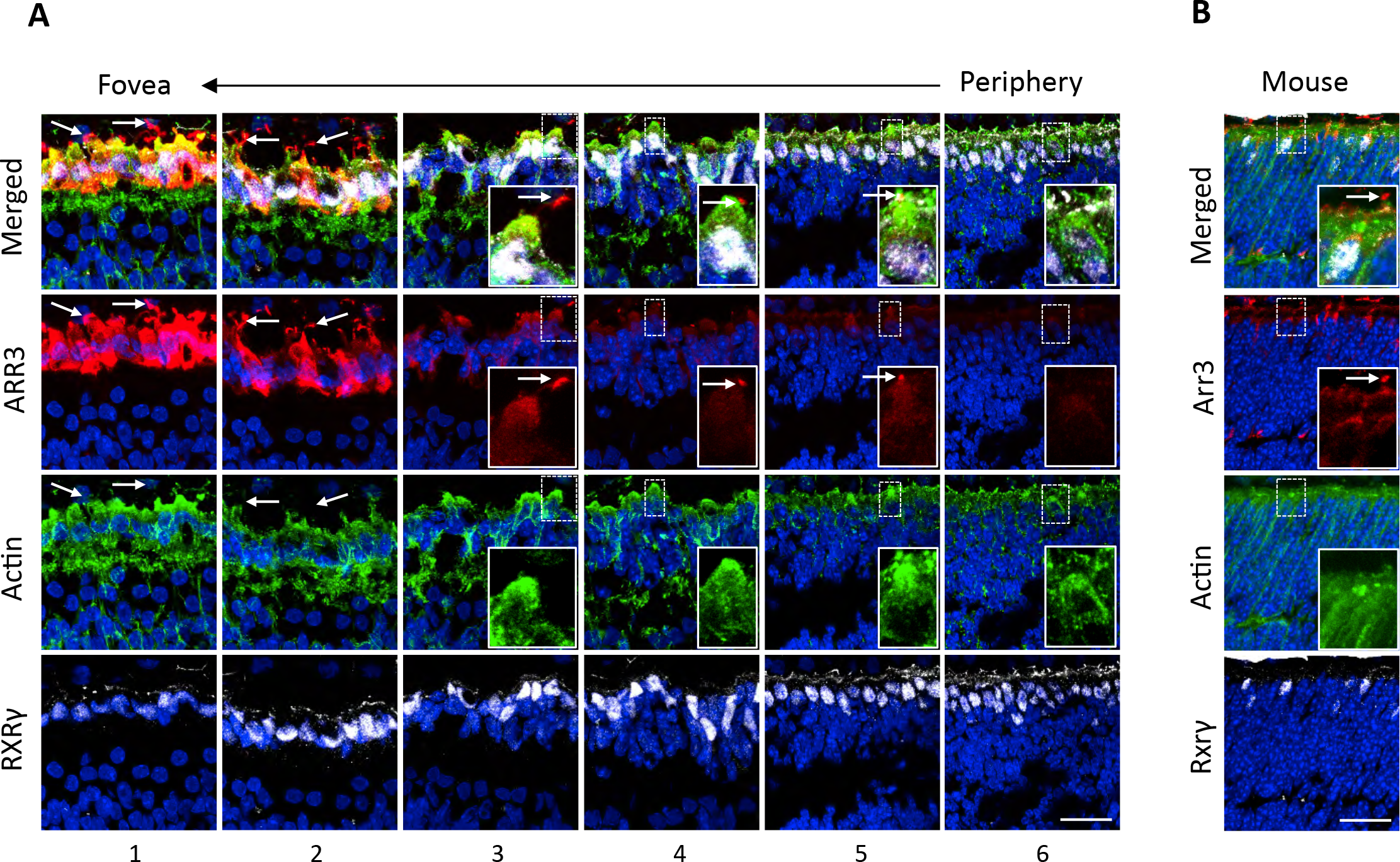
ARR3 expression coincides with the emergence of cone precursor outer segments. **A.** Human week 20 retina co-stained for ARR3 (red), Actin (green), and RXRγ (white). Arrow represents the increasing maturation gradient from the retinal periphery (position 6) to the fovea (position 1). Concentrated actin filaments in the apical portion of inner segments are thought to have a role in the development of photoreceptor outer segments and calycal processes (1–3). These concentrated actin filaments were detected at the apical surface in all RXRγ+ cone precursors in central retina (positions 1-4), in a minority of cone precursors at the more peripheral position 5, and in rudimentary cone precursor structures that did not reach the apical surface at the most peripheral position 6. RXRγ+ cone precursors with strong apical actin staining (at positions 4 and 5) had ~0.5 micron diameter outer segment buds that stained strongly for ARR3 (arrows), whereas cone precursors that lacked apical actin staining (at positions 5 and 6) lacked ARR3+ buds. **B**. Mouse P6 retina co-stained for Arr3, Actin, and Rxrγ. At P6, Arr3 was detected in Rxrγ+ cone precursor cell bodies and in ~0.5 micron outer segment buds (arrow) that first emerge at this age (3, 4) and were apical to highly actin-positive inner segments. Insets, enlarged view of actin clusters. Confocal images show maximum intensity projection of image stacks. Scale bars, 20 μm.

**Figure S2.**
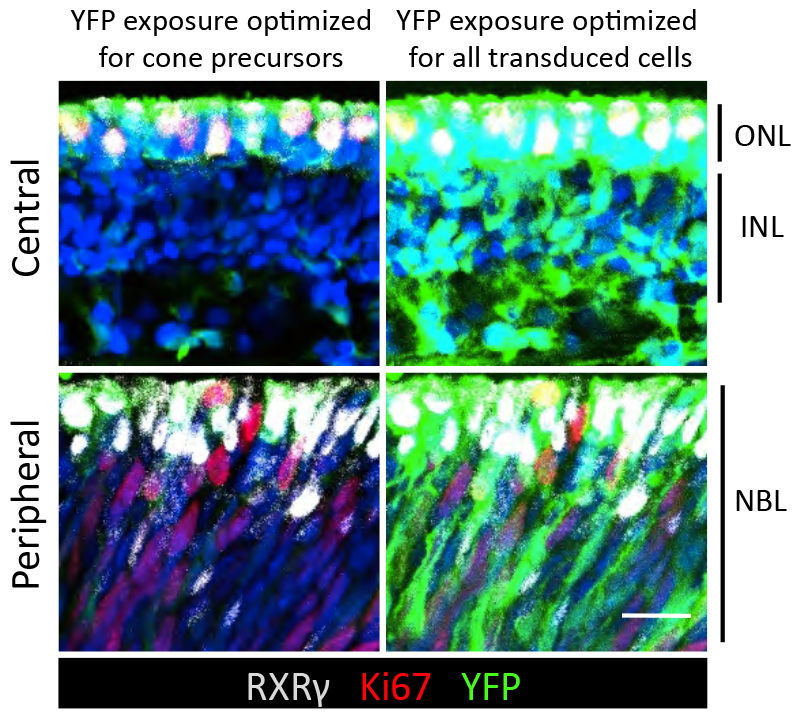
Lentiviral transduction of multiple layers of human retina. pLKO-YFP-sh*RB1* lentiviral transduction of cells throughout the inner nuclear layer (INL) and outer nuclear layer (ONL) of the central retina and throughout the neuroblastic layer (NBL) of the peripheral retina is evident with imaging optimized to detect all transduced cells. Cone precursors in the ONL shows brightest YFP signal and were best resolved with exposures optimized for this cell type. Scale bar, 20 μm.

**Figure S3.**
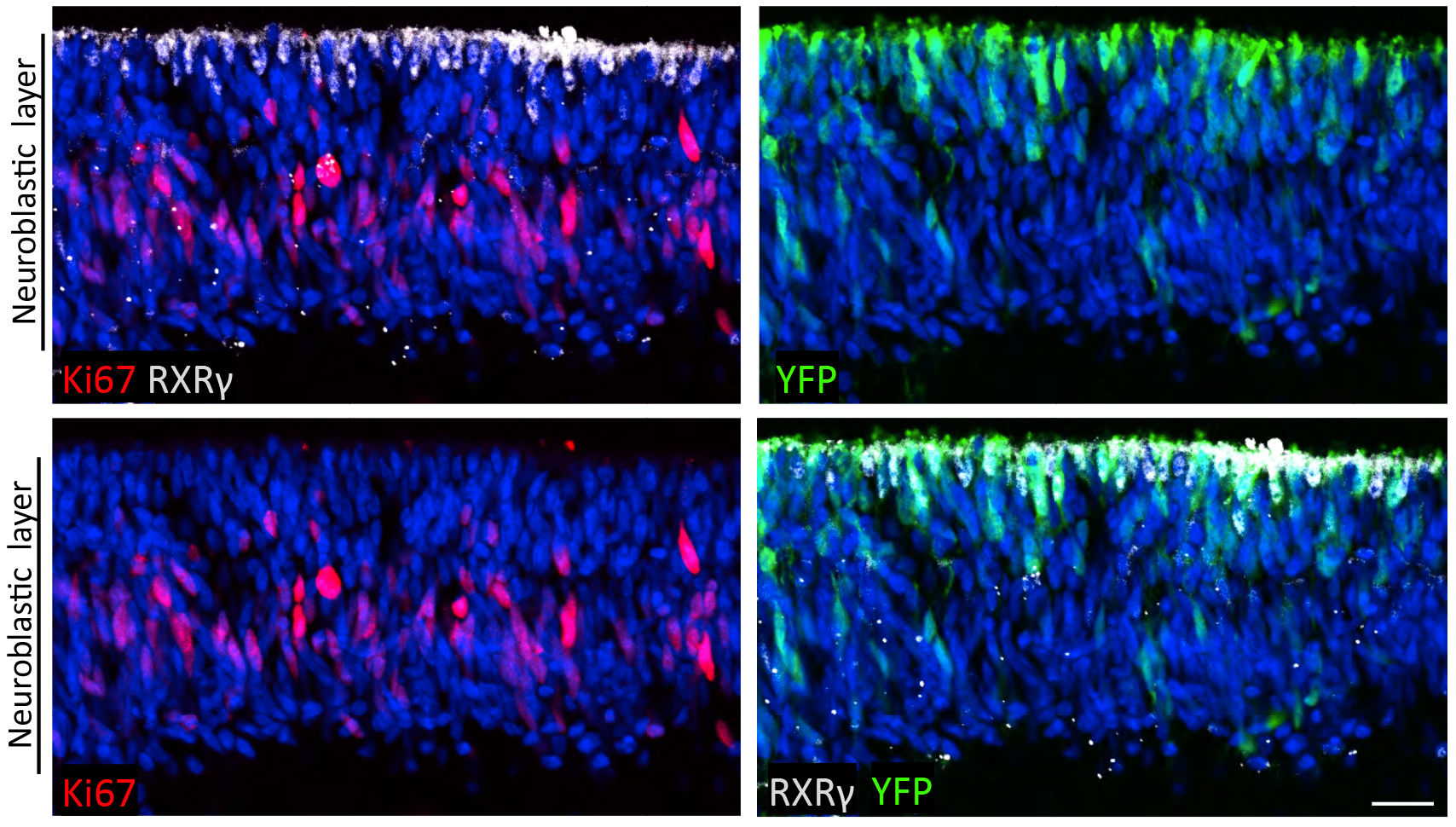
Proliferating RXRγ(-) cells in sh*SCR*-transduced human retina. sh*SCR* transduced retina at 12 DIC shows Ki67+,RXRγ(-) cells in the neuroblasti’c layer, similar to that observed in peripheral *shRbl*-transduced retina (Fig. 1C-E). Scale bar, 20μm.

**Figure S4.**
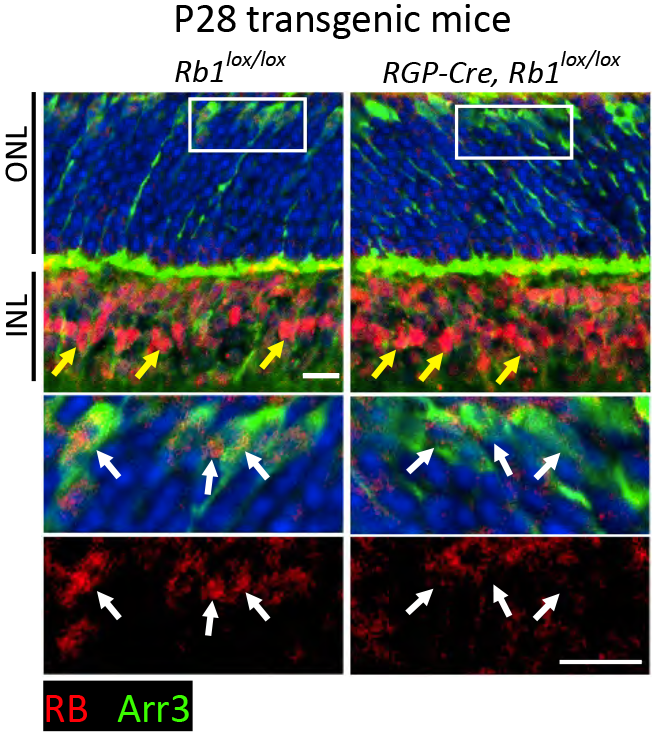
Cone-specific *Rb1* knockout in *RGP-Cre;Rb1*^*lox/lox*^ mouse retina at postnatal day P28. Upper panels show sections traversing the outer nuclear layer (ONL) and inner nuclear layer (INL). Boxed ONL regions are enlarged in lower panels. White arrows show Arr3+ cones and yellow arrows show Müller glia. Residual signal in the P28 *RGP-Cre*;*Rb1*^*lox/lox*^ ONL is intercellular and likely background. `Scale bar, 10μm`

**Figure S5.**
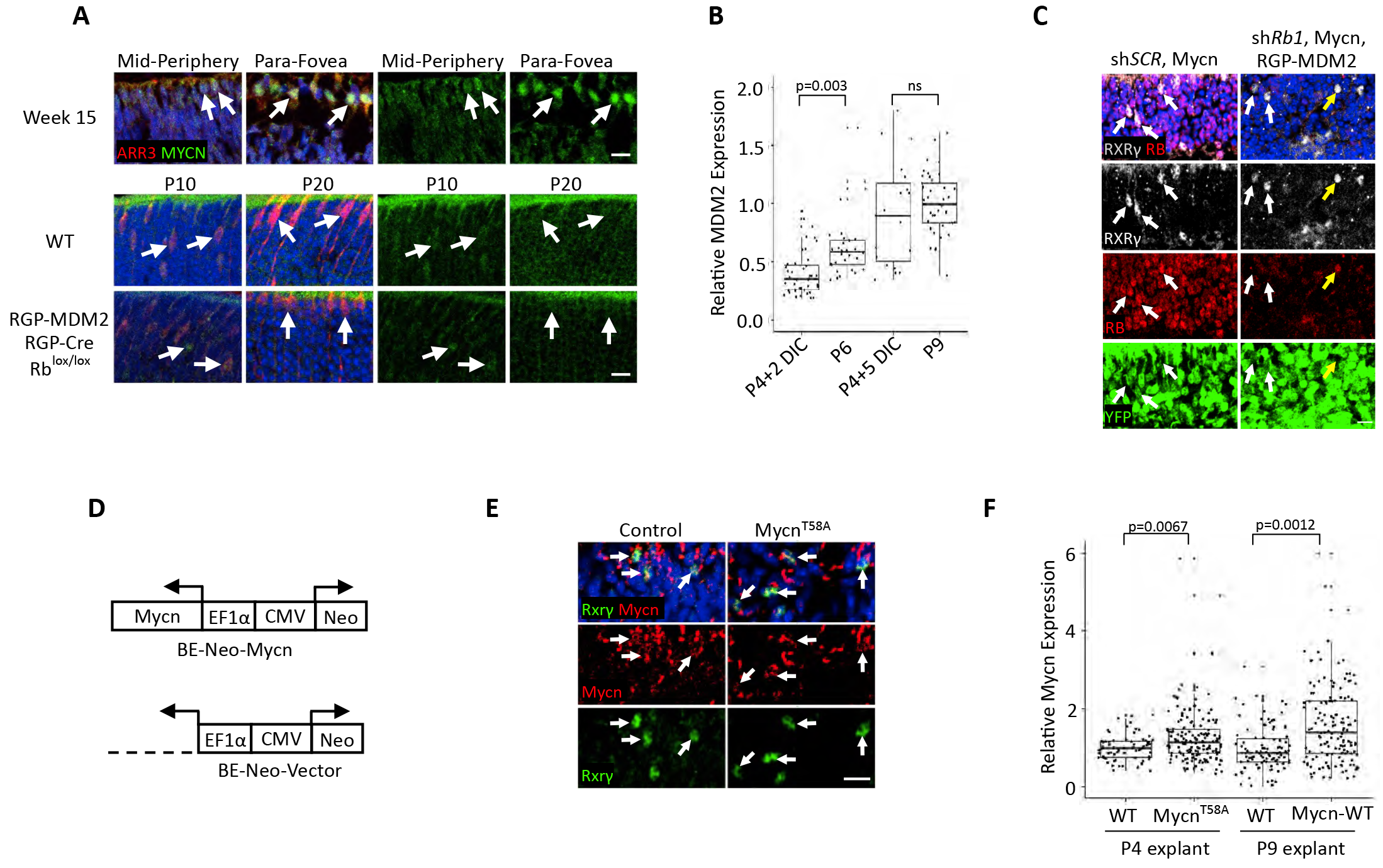
MDM2 and Mycn overexpression in transgenic and lentivirus-transduced mouse retinae. **A.** Developmental increase in MYCN expression in human but not in mouse cone precursors. *Upper panels*, ARR3+ cells in human week 15 retinae show increasing MYCN expression from the mid-periphery to parafovea. *Lower panels*, Mycn was weakly expressed in Arr3+ cells at P10 and not detected above background levels at P20 in both WT and *RGP-MDM2;RGP-Cre;Rb*^*lox/lox*^ mice retinae demonstrating that the MDM2 transgene did not alter endogenous Mycn expression. **B.** Quantitation of MDM2 expression in Rxrγ+ cone precursors in cultured vs. *in vivo* developed *RGP-MDM2* littermate retinae. MDM2 levels were compared between P6 retina and P4 retinal explants cultured for 2 days or between P9 retinae and P4 explants cultured for 5 days. Each dot represents a quantitatively imaged cell outlined by Rxrγ signal, and box plots show median (line inside box), upper and lower quartiles (box borders), and data range (whiskers). Significance was assessed by t-test. P values are as shown; ns, not significant. **C.** *Rb1* KD in a P4+7 DIC retinal explant co-stained for Rxrg, Rb, and YFP. White arrows, infected (YFP+) cells that are Rb+ after transduction with sh*SCR* but not with sh*Rb1*. Yellow arrow, uninfected (YFP-) cell with Rb expression. **D.** Lentiviral constructs for transduction of intact murine retina with ectopic Mycn or Mycn^T58A^ (BE-Neo-Mycn^(T58A)^) or with the empty vector (BE-Neo). **E.** Mycn expression in Rxrγ+ cells of P4 retinal explants transduced with Mycn^T58A^ at 7 DIC. **F.** Quantitation of Mycn levels in Rxrγ+ cells (*left*) or Arr3+ (*right*) at 7 DIC in P4 and P9 explants, respectively. Box plots and significance tests are as in panel B. Scale bars in A, C, and E, 10 μm.

**Figure S6.**
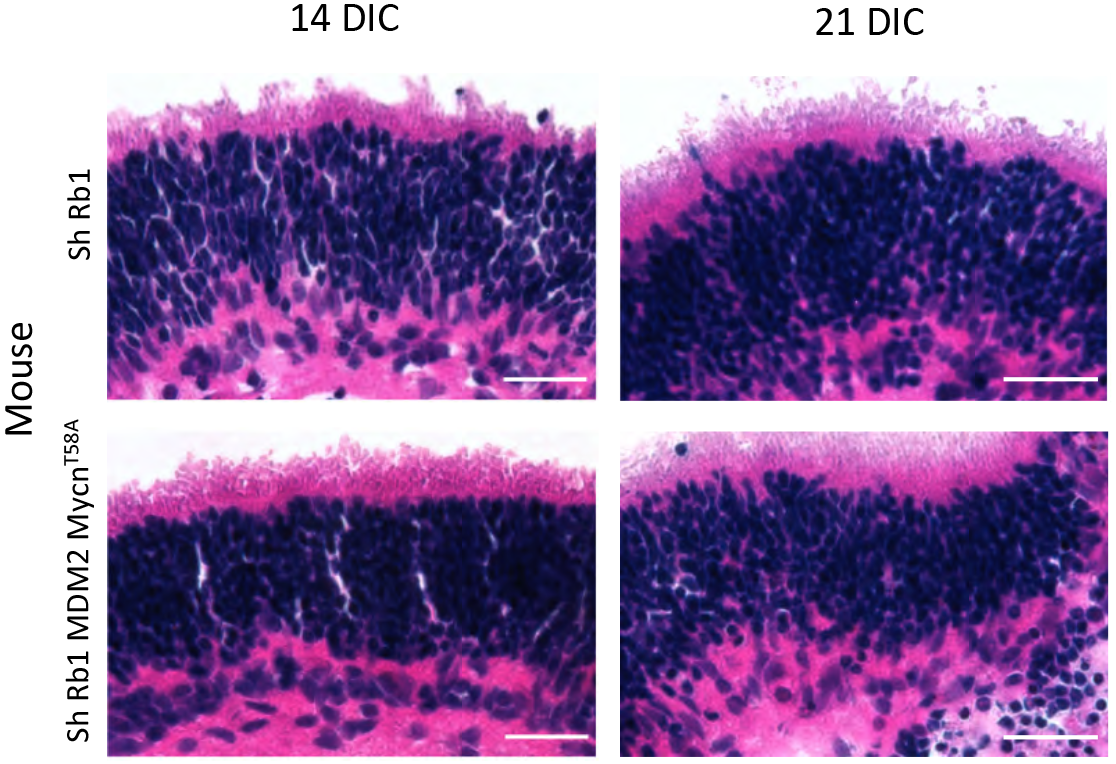
Absence of rosettes and fleurettes in Rb-depleted cultured murine retinae. Hematoxylin & eosin staining of explanted P4 wild type retina at 14 and 21 days post-Rb KD (*top*) and explanted P4 *RGP-MDM2* retina at 14 and 21 days post-Rb KD and Mycn overexpression (*bottom*). Scale bars, 20 μm.

**Table S1.**
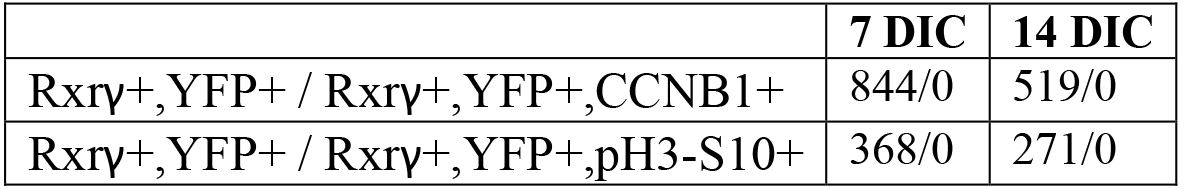
Lack of G2-M markers in *RGP-MDM2* cone precursors after Rb KD and Mycn overexpression.

|  | 7 DIC | 14 DIC |
| --- | --- | --- |
| Rxrγ+, YFP+ / Rxrγ+, YFP+, CCNB1+ | 844/0 | 519/0 |
| Rxrγ+, YFP+ / Rxrγ+, YFP+, pH3-S10+ | 368/0 | 271/0 |

**Table S2.** Antibodies used in this study.

| Name | Clone | Species | Source | Catalog number | Titre |
| --- | --- | --- | --- | --- | --- |
| Actin | - | Goat | Santa Cruz | SC-1616 | 1:100 |
| ARR3 (LUMif-hCAR) | - | Rabbit | Cheryl Craft (5) |  | 1:5000 |
| ARR3 | - | Rabbit | Millipore | AB15282 | 1:500 |
| CCNB1 | GNS1 | Mouse | Santa Cruz | SC-245 | 1:100 |
| GFP | - | Goat | Abcam | ab6673 | 1:500 |
| Ki67 | B56 | Mouse | BD Biosciences | 550609 | 1:200 |
| L/M-Opsin | - | Rabbit | Millipore | AB5405 | 1:500 |
| MDM2 | IF2 | Mouse | Calbiochem | OP46 | 1:100 |
| MDM2 | SMP14 | Mouse | Santa Cruz | SC-965 | 1:150 |
| MYCN | NCM II-100 | Mouse | Santa Cruz | SC-56729 | 1:100 |
| p130 | 10/Rb2 | Mouse | BD Biosciences | 610261 | 1:100 |
| p16 | - | Rabbit | Proteintech | 10883-1-AP | 1:100 |
| PH3 | - | Rabbit | Millipore | ab5176 | 1:200 |
| RB | G3-245 | Mouse | BD Biosciences | 554136 | 1:100 |
| RXRγ | - | Rabbit | Santa Cruz | sc-555 | 1:1000 |
| RXRγ | G-6 | Mouse | Santa Cruz | sc-514134 | 1:200 |

